# Biological subtyping of autism via cross-species fMRI

**DOI:** 10.1101/2025.03.04.641400

**Authors:** Marco Pagani, Valerio Zerbi, Silvia Gini, Filomena Alvino, Abhishek Banerjee, Andrea Barberis, M. Albert Basson, Yuri Bozzi, Alberto Galbusera, Jacob Ellegood, Michela Fagiolini, Jason Lerch, Michela Matteoli, Caterina Montani, Davide Pozzi, Giovanni Provenzano, Maria Luisa Scattoni, Nicole Wenderoth, Ting Xu, Michael Lombardo, Michael P Milham, Adriana Di Martino, Alessandro Gozzi

## Abstract

It is frequently assumed that the phenotypic heterogeneity in autism spectrum disorder reflects underlying pathobiological variation. However, direct evidence in support of this hypothesis is lacking. Here, we leverage cross-species functional neuroimaging to examine whether variability in brain functional connectivity reflects distinct biological mechanisms. We find that fMRI connectivity alterations in 20 distinct mouse models of autism (n=549 individual mice) can be clustered into two prominent hypo- and hyperconnectivity subtypes. We show that these connectivity profiles are linked to distinct signaling pathways, with hypoconnectivity being associated with synaptic dysfunction, and hyperconnectivity reflecting transcriptional and immune-related alterations. Extending these findings to humans, we identify analogous hypo- and hyperconnectivity subtypes in a large, multicenter resting state fMRI dataset of n=940 autistic and n=1036 neurotypical individuals. Remarkably, hypo- and hyperconnectivity autism subtypes are replicable across independent cohorts (accounting for 25.1% of all autism data), exhibit distinct functional network architecture, are behaviorally dissociable, and recapitulate synaptic and immune mechanisms identified in corresponding mouse subtypes. Our cross-species investigation, thus, decodes the heterogeneity of fMRI connectivity in autism into distinct pathway-specific etiologies, offering a new empirical framework for targeted subtyping of autism.

## Introduction

Autism spectrum disorder (ASD; hereafter referred to as autism) is characterized by highly heterogeneous phenotypic presentation, encompassing variable expression of core diagnostic symptoms and associated features, such as language, intellectual, motor, and adaptive functioning^1–3^. This diversity also manifests in multiple neuroimaging endophenotypes, including differences in brain activation patterns, functional connectivity, and morphometric features^3–8^. Recent investigations have revealed a similarly striking heterogeneity in the genetic and biological processes known to be associated with autism^9–11^. Specifically, large-scale genetic studies have shown that the high heritability of autism involves multiple (>100) rare, highly penetrant mutations, as well as common genetic risk variants ^9–11^. These genetic factors, alone or in combination, affect highly heterogeneous biological pathways, including synaptic activity, neurogenesis, cell migration and gene transcription ^12,13^. Beyond genetics, environmental influences - particularly prenatal inflammatory conditions and immune dysfunction - have also been shown to modulate autism risk^14^.

A common assumption exists that the phenotypic heterogeneity observed in autism directly reflects underlying pathobiological heterogeneity^15–18^. However, causal evidence in support of this hypothesis is lacking. Current genetic analyses do not allow for a reliable biological stratification of autism, since only 20% of individuals harbor clinically pathogenic rare variants, and no single mutation accounts for more than 1% of cases^11^. Given these limitations, recent efforts have focused on identifying phenotypically defined subtypes within autistic population. Statistical clustering of clinical and neuroimaging phenotypes has been used to identify putative autism subtypes, i.e., subgroups of autistic individuals characterized by more uniform clinical and/or neural phenotypes, putatively representing different pathobiological mechanisms^16–19^. However, despite the potential of this approach, direct evidence of its validity remains elusive. Most subtyping studies to date lack plausible neurobiological validation, and rely, at best, on tentative associations between neuroimaging metrics and normative gene expression patterns^15,18^.

Cross-species approaches enabling the biological decoding of autism-relevant phenotypes^20^ could bridge this critical knowledge gap, offering a pivotal avenue for advancing autism research. Rodent models offer a unique experimental approach to isolate and probe the effect of autism-relevant etiological factors on brain connectivity, with minimal genetic or environmental confounds^20,21^. Leveraging technical advances in cross-species functional neuroimaging, we and others have highlighted remarkably conserved fMRI connectivity alterations in both clinical populations and mouse lines harboring corresponding autism-relevant genetic variants^22–25^. Large-scale functional neuroimaging across multiple mouse autism models, thus, offers an unprecedented opportunity to biologically decode autism heterogeneity into etiologically distinct dysconnectivity signatures, and to potentially guide cross-species subtyping. Within this translational framework, rodent models have proven instrumental in modeling key autism-relevant immune alterations, as well as many of the multiple high-penetrance, rare genetic variants that constitute approximately 20% of autism’s genetic architecture ^26^. Importantly, postmortem studies have highlighted substantial overlap between pathways dysregulated in idiopathic autism, and those affected by rare genetic variants, further validating the use of rodent models to probe autism-relevant pathobiological mechanisms^13,27^.

Building on this notion, we reason that cross-species fMRI map decoding^20,25,28^ could empirically guide the identification of brain dysconnectivity subtypes in autism reflecting some of the biological pathways modelled in rodents. Using this approach, here, we show that variability in brain functional connectivity encodes dissociable biological pathways. Specifically, we report that fMRI connectivity alterations in 20 mouse models of autism can be clustered into two dominant hypo- and hyperconnectivity subtypes reflecting synaptic and transcriptional/immune-related pathways, respectively. Guided by these rodent findings, we identify analogous hypo- and hyperconnectivity subtypes in fMRI scans from autistic individuals and link these patterns to autism-relevant synaptic and immune mechanisms. Our work documents that heterogeneous fMRI connectivity in idiopathic autism encodes for dissociable pathobiological mechanisms, laying a foundation for biologically informed clinical subtyping of the autism spectrum.

## Results

### fMRI dysconnectivity in 20 autism mouse models clusters into dominant hypo- and hyperconnectivity subtypes

Previous investigations using resting-state fMRI have revealed highly heterogeneous patterns of atypical connectivity (here termed fMRI dysconnectivity) in autistic individuals ^29,30^. While this heterogeneity is often assumed to reflect underlying etiopathological variation, direct evidence in support of this hypothesis is lacking. To empirically probe this notion, we examined resting state fMRI connectivity in an aggregated database of 20 different mouse lines modelling autism-relevant genetic mutations spanning different pathways (e.g., synaptic mechanisms, protein translation, transcriptional regulation, chromatin remodeling), as well as immune-related mechanisms known to be relevant to autism. We hypothesized that if biological heterogeneity significantly contributes to the phenotypic variability observed with fMRI, discrete dysconnectivity clusters should be associated with distinct etiological mechanisms as indicated by their gene mutations and the affected transcriptional pathways. By using a relatively large number of different mouse models, we aimed to capture the diverse etiological landscape of autism, enabling us to empirically test our hypothesis.

**Table 1** provides detailed information on the employed mouse model database, which represents an extension of a previously described dataset ^31^. Importantly, each scanned model contains a control group of wild-type control littermates. This allowed us to precisely probe the fMRI connectivity alterations characterizing different pathobiological mechanisms with negligible environmental and genetic confounds. To facilitate cross-species translation of mouse findings to human populations, connectivity differences associated with each etiology were mapped at the voxel level using weighted-degree centrality. This metric quantifies the mean fMRI connectivity of each voxel ^32^ and has previously revealed comparable brain dysconnectivity signatures in rodents and humans harboring syntenic autism-risk ^23–25^.

**Table 1.**
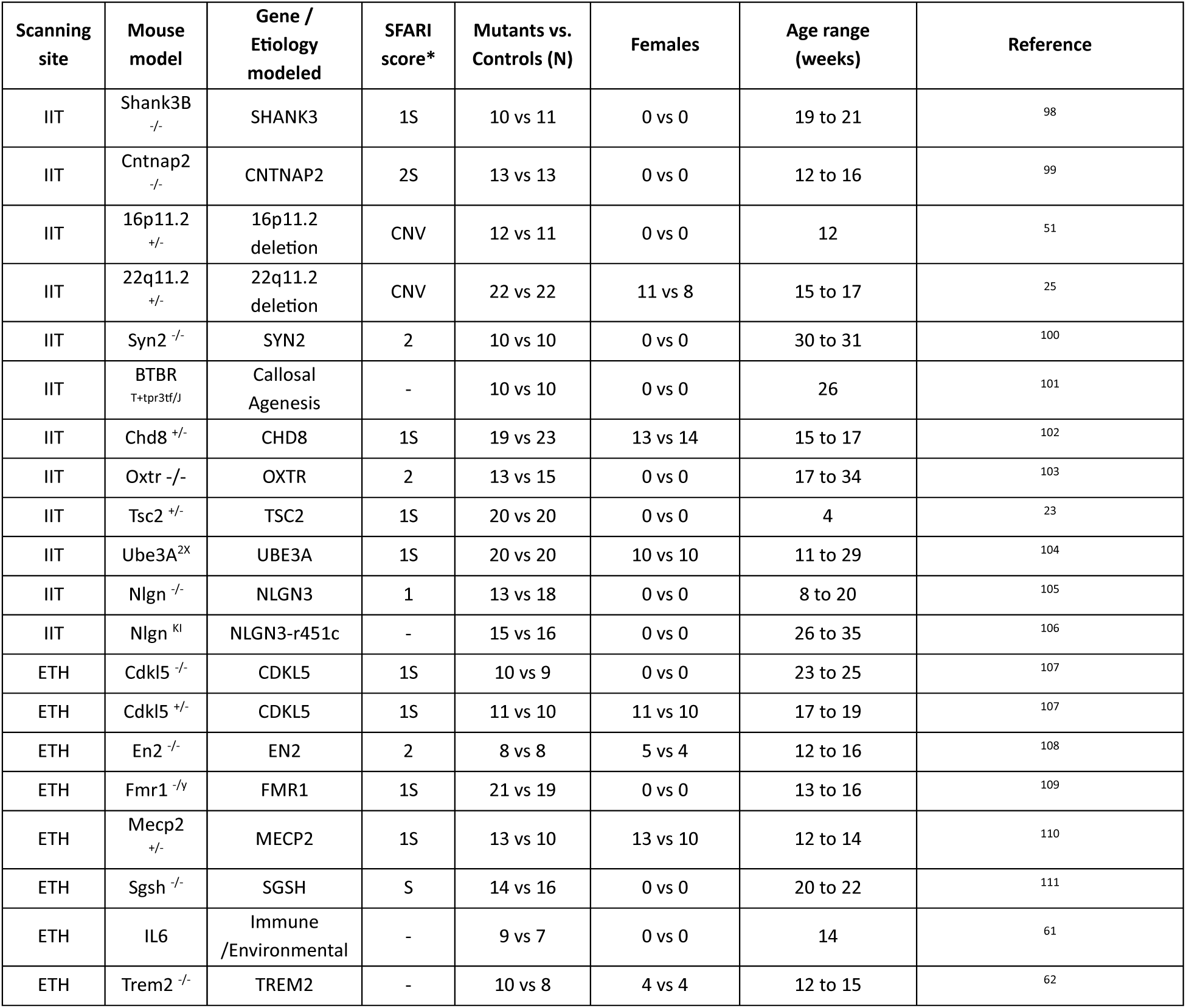
Autism-related mouse models. Scanning site: IIT, Istituto Italiano di Tecnologia, Rovereto, Italy. ETH, ETH Zurich, Switzerland. *SFARI scores were derived from https://gene.sfari.org/ in January 2025 and indicate the strength of the evidence of their implications in autism on a scale ranging from 1 (high) to 3(low). S: syndromic.

Voxel-wise quantification of fMRI dysconnectivity differences (i.e., mutant vs. wild-type) across the 20 autism models revealed a spectrum of dysconnectivity, ranging from marked hypoconnectivity (i.e., decreased fMRI connectivity in mutants) to marked hyperconnectivity (i.e., increased fMRI connectivity in mouse models, **Figure 1a,b**). This finding indicates that biological variability is a key determinant of connectivity heterogeneity in mouse models relevant to autism. Interestingly, while most models exhibited a combination of both hyper- and hypoconnected voxels, a clear polarization of the dysconnectivity landscape was also evident, with multiple mouse models exhibiting either predominant hypoconnectivity (e.g., En2, Shank3, 22q11.2, 16p11.2, Ube3A, SGSH) or predominant hyperconnectivity (e.g., Cdkl5[ko], fMRI1, Chd8, Tsc2, IL6; **Figure 1a,b**). In keeping with this notion, hierarchical clustering of fMRI dysconnectivity revealed two dominant subtypes characterized by prominent hypo- (n=11 mouse models) and hyperconnectivity (n=9, **Figure 1c**), respectively. While additional partitions could be identified at a lower hierarchical level (e.g., with k=3, see **Methods**), subsequent analyses focused on the two prominent subtypes. We reasoned that if the observed fMRI dysconnectivity reflects dissociable biological mechanisms, functionally divergent clusters like those identified here should be associated with distinct molecular profiles. This two-cluster approach also provided sufficient mouse models per subtype to generate robust (Cohen’s *d* >0.8) subtype-level dysconnectivity maps. These were obtained through a conjunction analysis of all fMRI dysconnectivity patterns, each corresponding to a specific mouse model, within each subtype **(Figure 1d)**. The resulting cross-etiological dysconnectivity maps represent brain regions that are most consistently vulnerable to hypo- or hyperconnectivity in each subtype.

**Figure 1.**
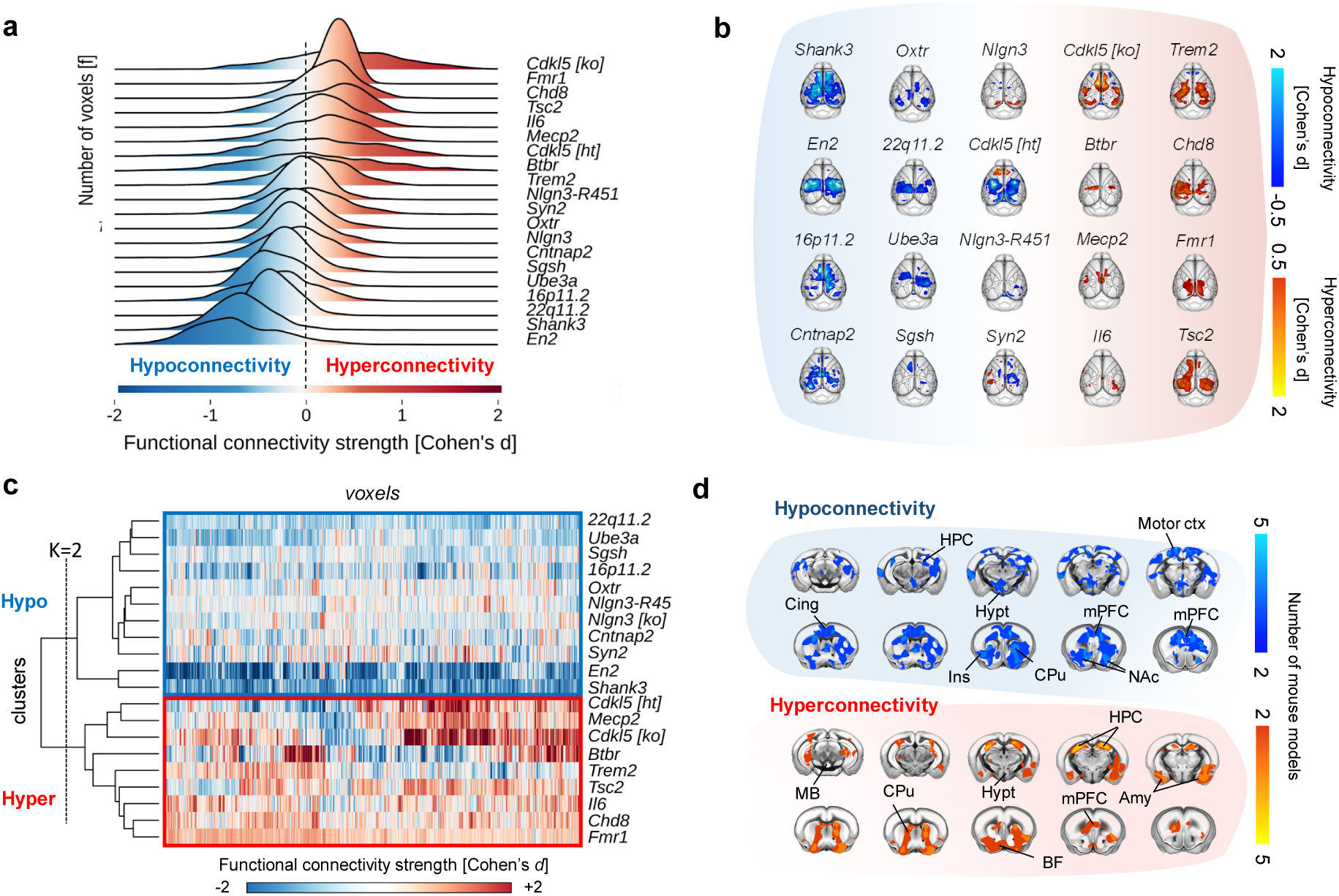
fMRI connectivity in 20 autism mouse models clusters into dominant hypo- and hyperconnectivity subtypes. **a)** Ridgeline plot quantifications of whole-brain fMRI connectivity strength differences (mutant vs. control) for the 20 autism-relevant mouse models examined. Ridgelines show voxel distribution of fMRI connectivity strength indexed by Cohen’s d color-coded by connectivity difference (blue: lower connectivity in mouse mutants vs. control, i.e., hypoconnectivity; red: higher connectivity in mouse mutants vs. control, i.e., hyperconnectivity). **b)** Top-view brain maps showing fMRI connectivity differences (with Cohen’s d > 0.5) in the 20 autism-related mouse models relative to wild-type littermates. Blue/light blue indicates hypoconnectivity, red/yellow indicates hyperconnectivity in mutants. **c)** Heatmap of Cohen’s d values across brain regions. Hierarchical clustering revealed two distinct connectivity subtypes: a hyperconnectivity cluster (n=9 mouse models) and a hypoconnectivity cluster (n=11 mouse models). Hypo, hypoconnectivity; Hyper, hyperconnectivity. **d)** Coronal brain views highlighting voxels with consistent hypo- or hyperconnectivity in the two fMRI dysconnectivity subtypes. Color intensity reflects the number of autism-related mouse models showing fMRI dysconnectivity (blue, hypoconnectivity, top panel; red/yellow, hyperconnectivity, bottom panel) in each subtype (threshold: Cohen’s d > 0.8). Amy, amygdala; BF, basal forebrain; Cing, anterior cingulate; CPu, caudoputamen; HPC, Hippocampus; Motor ctx, motor cortex; Hypt, hypothalamus; Ins, Insula; mPFC, medial prefrontal cortex; MB, mid brain; NAc, nucleus accumbens.

Interestingly, these subtype specific maps revealed both overlapping and distinct regional patterns. Some anatomical substrates, including the medial prefrontal cortex, striatum, and basal forebrain, were susceptible to either hypo- and hyperconnectivity across subtypes. Other regions showed instead subtype-specific alterations. For example, the hippocampus and amygdala were predominantly affected in the hyperconnectivity subtype, while the hypothalamus and somatomotor cortex were specifically altered in the hypoconnectivity group. These findings demonstrate that the dysconnectivity landscape across 20 autism mouse models segregates into two dominant, functionally opposed patterns. Importantly, our identification of the brain regions most affected in each subtype also provided a translational framework for comparing dysconnectivity signatures across species.

### fMRI hypo- and hyperconnectivity reflect dissociable biological pathways

The identification of dominant hypo- and hyperconnectivity subtypes in our mouse database enabled us to empirically investigate whether different patterns of autism-relevant dysconnectivity reflect dissociable pathobiological pathways. To test this hypothesis, we generated two aggregate sets of molecular pathways, each specifically associated with fMRI hypo- or hyperconnectivity. We began by constructing *in silico* two protein-protein mega-interactomes, each comprising the genetic and immune-related etiologies belonging to either the hypo- or hyper-connectivity subtypes, along with their interacting genes (**Figure 2a**). We next filtered out the genes present in both mega-interactomes, retaining only those uniquely represented in each subtype-specific gene set. This step allowed us to focus our subsequent investigations on pathways more likely unique to either the hypo- or hyperconnectivity subtypes. We finally applied a gene ontology analysis to identify, for each of the resulting interactomes, the prevalence of molecular pathways known to be dysregulated in autism ^33^.

**Figure 2.**
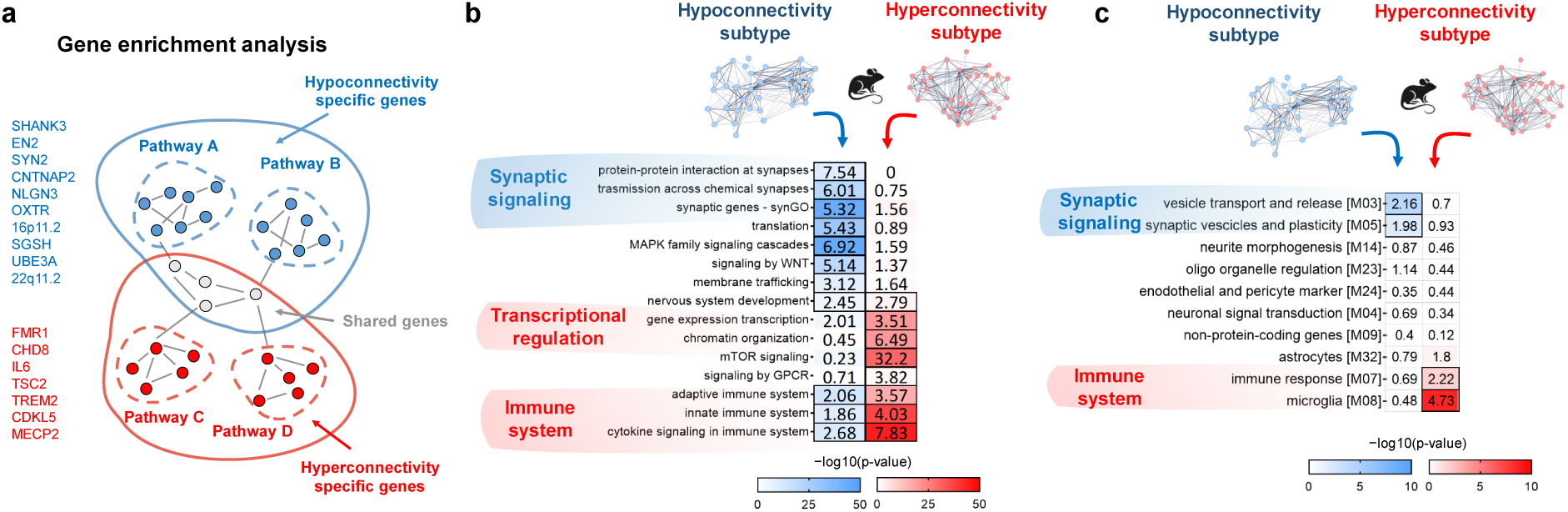
Distinct signaling pathways underlie fMRI connectivity subtypes in rodents. **a)** Illustrative schematic of gene enrichment analysis used to link autism relevant pathways to rodent hypo- and hyperconnectivity subtypes. Autism-risk genes and immune factors (i.e., IL-6) modelled in mouse lines associated with either subtype (listed in blue and red typeface, respectively) were used as seed genes to generate mouse line-specific protein-protein interactomes. Within each subtype, these interactomes were then concatenated, and upon removal of shared genes, we generated two subtype-specific gene interactomes. Filled circles represent individual genes, gray links indicate gene interactions. Dashed lines delineate individual interactomes and solid lines outline the two concatenated interactomes. **b)** Heatmap displaying significant enrichment for autism-relevant pathways in the two interactomes. **c)** Heatmap displaying significant enrichment for modules of genes differentially expressed in autism ^13^ for each of the two interactomes. The odds ratio for hypoconnectivity subtype are shown in the left column (blue coloring); those for the hyperconnectivity subtype in the right column (red coloring). Thick cell borders indicate that enrichment is significant at q(FDR) < 0.05. We report the list of genes belonging to each interactome and the pathways we probed in **Supplementary Table 2.**

Using this approach, we identified a composite set of autism-relevant molecular pathways linked to hypo- or hyperconnectivity. Notably, these pathways were distinctly dissociable (**Figure 2b)**. Specifically, the hypoconnectivity subtype exhibited prominent enrichment for multiple synaptic-related ontologies, including genes involved in protein-protein interaction at the synapse (odds ratio, OR=7.54, p_(FDR)_=10^-9^), transmission across chemical synapses (OR = 6.01, p_(FDR)_ = 10^-20^), and synaptic functioning (SynGO, OR=5.32, p_(FDR)_=10^-42^). This subtype was also significantly enriched for pleiotropic molecular effectors that are also involved in the control of synaptic plasticity, such as MAPK signaling (OR = 6.92, p_(FDR)_=10⁻⁴³), membrane trafficking (OR=3.12, p_(FDR)_=10⁻^8^), protein translation (OR = 5.43, p_(FDR)_=10⁻^29^), and WNT pathway activity (OR=5.14, p_(FDR)_=10⁻¹⁹) ^34,35^. Immune-related mechanisms, such as innate (OR=1.86, p_(FDR)_=10^-8^), adaptive (OR=2.06, p_(FDR)_=10^-13^), cytokine signaling (OR=2.68, p_(FDR)_=10^-11^) were instead only weakly represented in this subtype.

By contrast, no synaptic-specific mechanisms were enriched in the hyperconnectivity interactome (all synaptic ontologies, qFDR>0.05, except for the pleiotropic mTOR pathway, OR=32.2, p_(FDR)_=10^-42^). This interactome, however, presented robust enrichment for immune-related pathways, such as cytokine signaling (OR=7.83, p_(FDR)_=10^-39^), innate immune response (OR=4.03, p_(FDR)_=10^-33^), adaptive immune response (OR=3.57, p_(FDR)_=10^-14^), as well as transcriptional mechanisms (e.g., chromatin organization, OR=6.49, p_(FDR)_=10^-29^, gene expression transcription OR=3.51, p_(FDR)_=10^-29^, **Figure 2b**). Importantly, these enrichments appeared to be robust to interactome stringency, as we found them to be fully preserved after increasing the number of interacting genes up to n=500 for each genetic or immune-related etiology (**Supplementary Figure 1**). To assess the robustness of these results against different sets of gene ontologies, we repeated enrichment analyses using gene modules previously described to be dysregulated in post-mortem brains of autistic individuals ^13^. Consistent with our original findings, we found that the hypoconnectivity subtype was robustly enriched for modules containing genes involved in regulation of synaptic activity, such as vesicle transport and release (OR=2.16, p_(FDR)_=10^-8^) and synaptic vesicles and plasticity (OR=1.98, p_(FDR)_=10^-4^). The hyperconnectivity interactome was instead significantly enriched for gene modules involved in immune signaling, such as immune response (OR=2.2, p_(FDR)_=10^-3^) and microglia (OR=4.73, p_(FDR)_=10^-17^, **Figure 2c**). As a note, because the Gandal et al. (2022) dataset did not include modules unequivocally related to transcriptional regulation, replicability of this pathway could not be evaluated. Taken together, these results reveal a set of dissociable mechanisms leading to diverging fMRI dysconnectivity in autism, with synaptic dysfunction leading to fMRI hypoconnectivity, whereas immune and transcriptional dysregulation are more prominently associated with fMRI hyperconnectivity.

### Reproducible hypo- and hyperconnectivity subtypes can be identified in human data

As summarized in **Figure 3**, following the identification of the two dominant connectivity subtypes in the rodent database, we probed the relevance of these findings in humans. Specifically, leveraging the cross-species translatability of fMRI connectivity ^20,25^, we asked whether analogous hypo- and hyper-connectivity subtypes could be identified in a large database of fMRI scans acquired in individuals with idiopathic autism. This approach was followed by a gene decoding of the autism-related hypo- and hyperconnectivity maps obtained in the human data ^25^.

**Figure 3.**
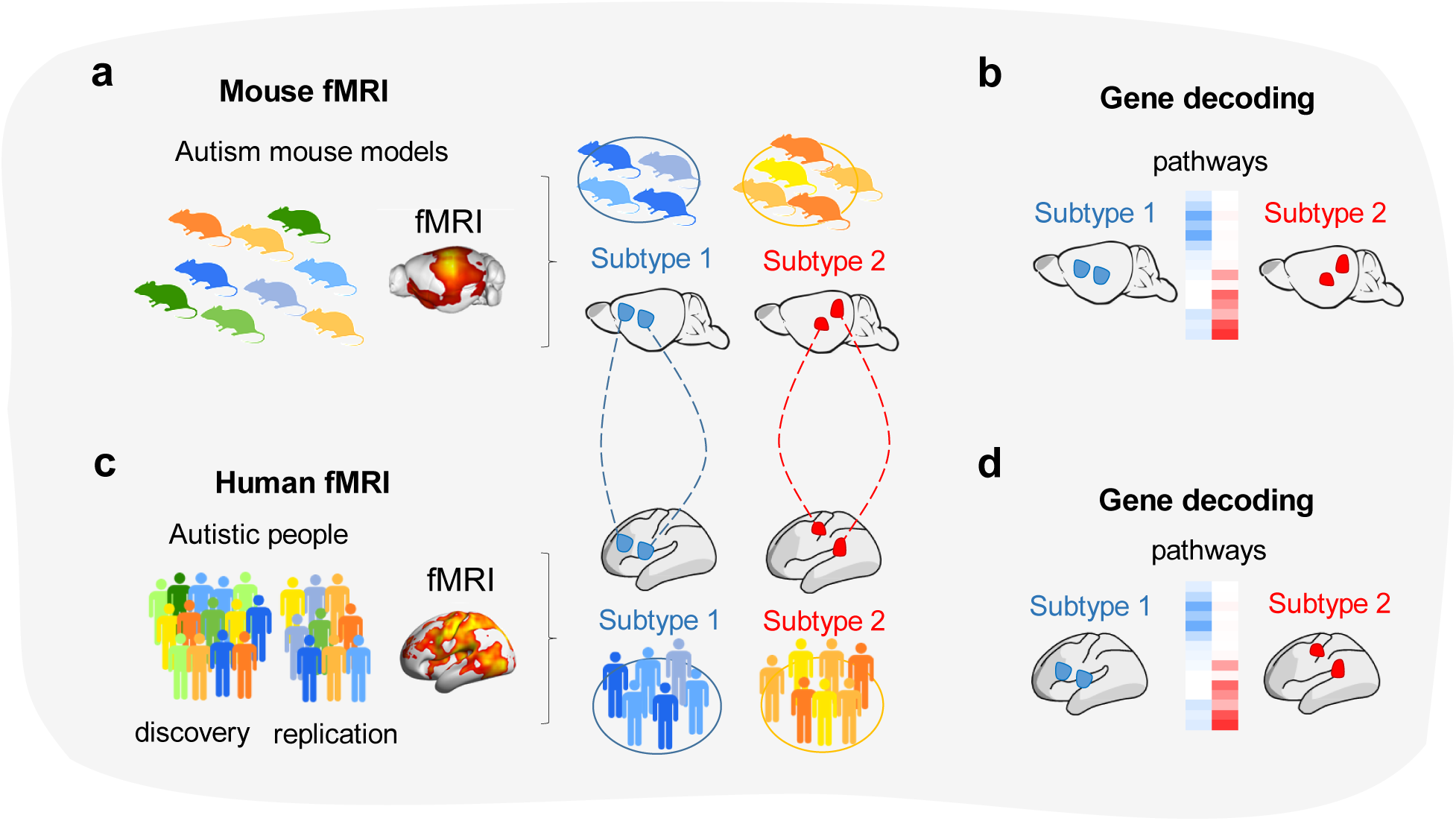
Cross-species identification of autism-related dysconnectivity subtypes. Schematic illustration of the workflow we used to identify hypo- and hyperconnectivity subtypes across species. **a)** We applied data-driven hierarchical clustering analyses to our rodent database to identify dominant hypo- and hyperconnectivity subtypes across autism mouse models relevant to autism. For each subtype we generated a dysconnectivity prior mask, consisting of a set of anatomical regions exhibiting hypo- or hyperconnectivity across models. **b)** Gene enrichment analyses were used to uncover molecular pathways associated with hypo- vs. hyperconnectivity in rodent autism models (**Supplementary Figure 1**). **c)** In humans, we computed fMRI dysconnectivity by comparing resting state fMRI data of autistic vs. neurotypical (NT) individuals. Leveraging the cross-species translatability of fMRI, we used a region-wise approach to quantify fMRI dysconnectivity in individuals with autism relative to NTs. Specifically, we selected fMRI scans exhibiting hypo- or hyperconnectivity in human brain regions corresponding to the dysconnectivity priors identified in rodents (**Supplementary Figure 2**). **d)** Finally, we performed gene enrichment analyses to investigate whether brain-decoded genes from each map were enriched for molecular ontologies or gene modules known to be associated with autism.

We examined an aggregated dataset of human low motion resting-state fMRI data, comprising n=940 individuals on the autism spectrum (age range 5–30 years old), and n=1036 age-matched neurotypical (NT) controls. The probed datasets comprised 38 data collections: 37 of them selected from the ABIDE repositories and one being a newly collected sample aggregated at CMI ^36–39^ (**Table 2)**. To evaluate subtype reproducibility, we *a priori* split this aggregated dataset into a discovery (78.5% of the aggregate sample, n=744 autistic individuals, n=807 NT controls) and a replication dataset (21.5% of the aggregate dataset, n=196 autistic individuals, n=229 NT controls; see **Methods**) matched for diagnosis, sex, age and in-scanner head motion.

**Table 2.** Breakdown of demographics and clinical scores by diagnostic group: autism and neurotypicals. For continuous variables, group mean, and standard deviations are reported in parentheses and minima and maxima are reported in brackets. **\***The data aggregate included 38 data collections, 18 from ABIDEI (Caltech, KKI, Leuven-1, Leuven-2, MaxMun, NYU, OHSU, Olin, Pitt, SDSU, Stanford, Trinity, UCLA-1, UCLA- 2, UM-1, UM-2, USM, Yale), 19 from ABIDEII (BNI-1, EMC-1, ETH-1, GU-1, IP-1, IU-1, KKI-1, KUL-3, NYU-1, NYU-2, OHSU-1, ONRC-2, SDSU-1, SU-2, TDC-1, UCD-1, UCLA-1, U-MIA-1, USM-1) and one from CMI; of them, 3 included only data from individuals on the autism spectrum (**Figure 4a**). **Full scale IQ was available in n=819 individual data in the autism diagnostic group and in n=968 of the NT group. ***Verbal IQ was available for n=709 individual data of the autism diagnostic group and of n=809 in the NT group. ****Non-Verbal NV IQ was available for n=726 individual data of the with autism group and for n=857 NT. ^Autism Diagnostic Observation Scale (ADOS) social affect (SA) calibrated severity scores (CSS) were available for n=520 individuals in the autism group. ^^Restricted and repetitive behaviors scores (RRB) were available for n=525 individuals in the autism group. ^^^ADOS total calibrated severity scores (CSS) score**s** were available for n=549 in the autism group (**Supplementary Methods).** ^^^^Number and percentage of individuals with autism and one or more **p**sychiatric cooccurring condition. Comorbidity was assessed in a subset of data with an autism spectrum disorder diagnostic label across 13 collections in n=436 individuals. ┼Number and percentage of individuals taking psychoactive medications among. Information on psychoactive medications was available for a subset of data (autism n=777 and NT n=822, across 27 collections). .FD, framewise displacement; r, Pearson correlation; χ2, chi-square statistics; t, unpaired t-test statistics.

|  | Autism | Neurotypical (NT) | Group comparisons, statistics and p-values |
| --- | --- | --- | --- |
| **Sample size**<br># | 940 | 1036 |  |
| **Data collections\***<br># | 38 | 35 |  |
| **Sex**<br>#M, #F | 811, 129 | 774, 262 | $\chi^2_{(1)} = 41.5$,<br>p<0.001 |
| **Age**<br>years | 13.7 (5.2) [5.1-29.2] | 14.1 (5.5) [5.9-29.9] | $t_{(1974)} = 1.93$,<br>p=0.06 |
| **Full Scale IQ\*\***<br>standard score | 105.9 (16.9) [41-149] | 113.1 (12.8) [71-151] | $t_{(1785)} = 10.3$,<br>p<0.001 |
| **Verbal IQ\*\*\***<br>standard score | 105.7 (17.8) [42-180] | 113.8 (13.7) [67-156] | $t_{(1516)} = 9.9$,<br>p<0.001 |
| **Non-Verbal IQ\*\*\*\***<br>standard score | 105.1 (17.3) [37-157] | 109.5 (13.7) [62-155] | $t_{(1581)} = 5.70$,<br>p<0.001 |
| **ADOS SA^**<br>CSS | 6.8 (2.1) [1-10] | - | - |
| **ADOS RRB^^**<br>CSS | 7.1 (2.5) [1-10] | - | - |
| **ADOS Total^^^**<br>CSS | 6.8 (2.2) [1-10] | - | - |
| **median FD**<br>mm | 0.067 (0.04) [0.01-0.19] | 0.062 (0.03) [0.01-0.19] | $t_{(1974)} = 2.97$,<br>p=0.003 |
| **Psychiatric comorbidity in autism^^^^**<br>n (%) | 241 (55.3%) | - | - |
| **Psychoactive medication use +**<br>n (%) | 246 (32%) | 5 (0.7%) | $\chi^2_{(1)} = 291.0$,<br>p<0.0001 |

To enable a cross-species extrapolation of our mouse results, we used a regional decoding approach focusing on evolutionarily conserved brain regions identified in the conjunction dysconnectivity maps of corresponding hypo- and hyperconnectivity rodent subtypes (**Methods, Supplementary Figure 2**). This enabled us to identify two subgroups (i.e., subtypes) of autistic individuals **(Figure 4a-d**), each showing robust (Cohen’s *d*>0.8) differences in fMRI connectivity relative to NT controls, consistent with hypo- and hyperconnectivity subtypes observed in the rodent dataset. Together, the two subtypes accounted for 24.1% of the discovery autism data (n=55, 7.4%; and n=124, 16.7%, for the hypo- and hyperconnectivity subtype, respectively). Quantifications of fMRI dysconnectivity confirmed that the evolutionary conserved regions showing significant hypo- or hyperconnectivity in humans were consistent with those we identified in corresponding rodent subtypes, thus validating our cross-species translation (**Supplementary Figure 3**). To probe the replicability of these findings, we repeated our cross-species decoding on a replication dataset identified *a priori*. Analyses of this dataset revealed two hypo- and hyperconnectivity subtypes, accounting for 29.1% of the autism replication data (n=19, 9.7% and n=38, 19.4% for the hypo- and hyperconnectivity subtype, respectively; **Figure 4b-c**). The topography and connectivity profile of the two subtypes were highly reproducible across the discovery and replication subtypes (hypoconnectivity subtype, Dice coefficient=0.74, r=0.67; hyperconnectivity subtype, Dice coefficient=0.96, r=0.73, **Supplementary Figure 4**). Collectively the identified subtypes accounted for 25.1% of the aggregated autism data examined (hypoconnectivity, N=74, 7.9%; hyperconnectivity, N=162, 17.2%, upon aggregation of discovery and replication autism datasets, i.e., n = 940). Across the combined sample, scans representing the hyperconnectivity subtype were present in all the data collections, while scans exhibiting hypoconnectivity were present in all but 10 data collections **(Figure 4d)**. To further corroborate the generalizability of these results, we repeated our cross-species decoding upon removal of the five largest data collections of autistic participants’ data. Results were largely consistent with our primary findings (**Methods** and **Supplementary Figure 5);** this suggests that our subtyping was not biased by the largest data collections.

**Figure 4.**
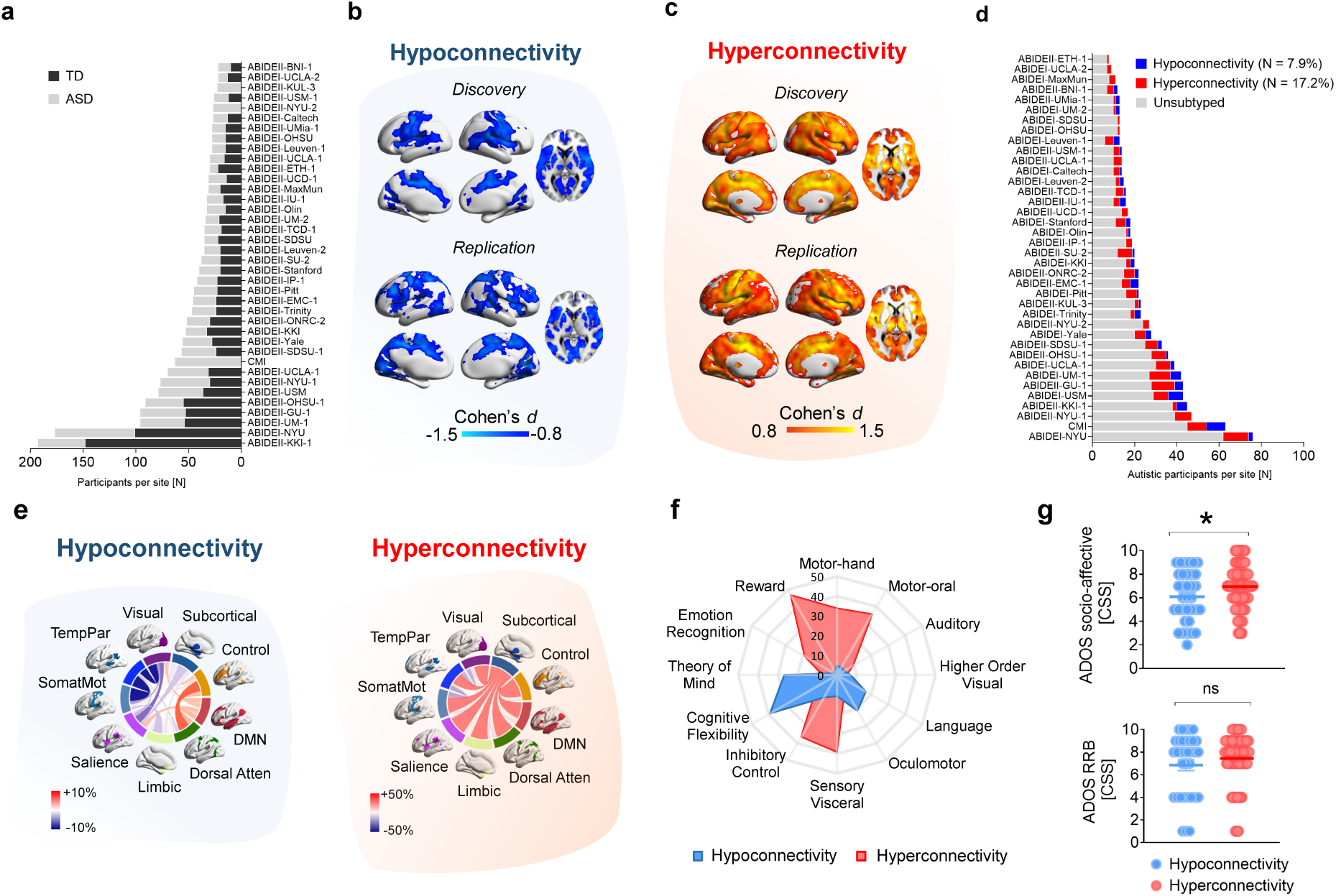
Replicable hypo- and hyperconnectivity subtypes can be identified in autism. **a)** Sample size distribution by data collection (ASD; light gray, NT; dark gray); ASD total n=940; NT total n=1036, n=38 data collections across 23 sites (**Table 2**). **b)** Hypoconnectivity subtype maps in discovery (top subpanel: n=55, 7.4%) and replication (bottom subpanel: n=19, 9.7%) datasets. Blue indicates regions exhibiting fMRI hypoconnectivity compared to NTs (Cohen’s d≤0.8). **c)** Hyperconnectivity subtype maps in discovery (top subpanel: n=124, 16.7%) and replication (bottom subpanel: n=38, 19.4%) datasets. Red/yellow indicates fMRI hyperconnectivity (Cohen’s d≥0.8). **d)** Distribution of autistic participants included in the hypo- or hyperconnectivity subtypes in the aggregated (discovery plus replication) autism dataset; Blue: hypoconnectivity subtype, n=74, across n=28 data collections; red: hyperconnectivity, n=162 across n=38 data collections), as well as those not subtyped (light gray n=704, across all 38 collections). **e)** Connectograms showing atypical fMRI network structure in hypo- (left) and hyperconnectivity (right) subtypes (upon regression of mean fMRI connectivity across 414 parcellation units) ^43,44^. Link thickness is proportional to the number of between-network edges displaying a significant difference in ASD vs. NT (red: increased, blue: decreased connectivity; t > 3.1, NBS corrected ^45^ at p < 0.05). **f)** Radar plot showing the percentage of overlap (range: 0–50%) between subtype mean regressed connectivity difference maps and 12 neuro-cognitive ontology probability maps ^46^. **g)** Autism subdomain severity scores (top subpanel: social affect (SA); bottom subpanel: restricted repetitive behaviors (RRB)) based on Autism Diagnostic Observation Schedule (ADOS; see Supplementary Methods) for each subtype. SA: hypoconnectivity subtype n=33, mean=6.1±2.2; hyperconnectivity, n=84, mean=7.0±1.8; t_(115)_=2.37, p_uncorr_=0.019, p_(FDR)_=0.030. RRB: hypoconnectivity subtype n=34, mean=6.9±2.8; hyperconnectivity, n=85, mean=7.5±2.2; t_(117)_=1.20, p_uncorr_=0.23, p_(FDR)_=0.23. Error bars: SEM; *p < 0.05; ns: non-significant; CSS: calibrated severity score. DMN, default mode network; Dorsal Atten, dorsal attentional network; Limbic, limbic network; Salience, salience network; SomatMot, somatomotor network; TempPar, temporoparietal network; Visual, visual network. NBS, network-based statistics.

### Hypo- and hyperconnectivity subtypes exhibit different network structure and are behaviorally dissociable

We further characterized the human subtypes in regard to their network structure, associated cognitive ontology maps ^40^, and autism symptomatology. To increase statistical power, analyses were carried out on the aggregate dataset combining the subtyped discovery and replication cohorts. Network comparisons of autism vs. NT data within subtypes (**Methods**) revealed specific network alterations. Markedly increased subcortico-cortical connectivity (t>3.1, p<0.05; **Figure 4e**) and reduced cortico-cortical connectivity between temporoparietal, visual, and somatomotor networks were noted in the hyperconnectivity subtype. In this subtype, the largest differences included increased connectivity within subcortical regions (53% of node-to-node links) and their connections to the salience network (44% of the 414 cortical and subcortical parcellation units ^41,42^). Conversely, the hypoconnectivity subtype displayed weaker, but significant network-level differences (left panel, t>3.1, p<0.05), particularly involving decreased connectivity between the somatomotor and temporoparietal networks (5% of links). Supporting a distinct network organization for the two subtypes, their connectivity matrices showed negligible spatial overlap (r=0.07, p=0.53). These findings suggest that the identified hypo- and hyperconnectivity patterns represent two functionally distinct subtypes characterized by different underlying brain network architecture. Consistent with this notion, reverse inference mapping ^40^ revealed that the network structure of these two subtypes is associated with distinct ontology cognitive maps. Specifically, the dysconnectivity map of the hypoconnectivity subtype overlapped with oculomotor, language, and cognitive areas, while the hyperconnectivity map was associated with sensory, visceral, motor, reward, and inhibitory control areas (**Figure 4f**).

Finally, to assess the behavioral profile of the two autism-related subtypes, we compared autism symptom severity using ADOS-2-based severity total and subtotal scores available in subsets of individuals (SA, n=117, RRB, n=119; CSS total, n=125); **Methods** and **Supplementary Methods**). Interestingly, on average, those in the hyperconnectivity subtype exhibited moderately increased total calibrated severity scores relative to individuals in the hypoconnectivity subtype (hypoconnectivity subtype n=38, mean=6.1±2.5; hyperconnectivity, n=87, mean=7.1±1.9; t_(123)_=2.20, p_uncorr_=0.020, p_(FDR)_=0.030). As shown in **Figure 4g**, comparisons of symptom subdomain severity scores revealed subtype differences for SA but not for RRB scores. Exploratory analyses showed no statistically significant differences between subtypes in regard to other available phenotypic data including age, IQ, sex, comorbidities rates and medication status (**Supplementary Table 1**). Taken together, these findings indicate that the identified hypo- and hyperconnectivity subtypes exhibit distinct functional network patterns and autism symptom severity.

### Hypo- and hyperconnectivity in humans recapitulate molecular pathways identified in rodent models

Given that similar connectivity subtypes exist in individuals with autism, we investigated whether they also reflect analogous molecular mechanisms. Using spatial gene decoding ^47^, we identified two sets of subtype-specific genes that are spatially enriched in the identified hypo- and hyperconnectivity subtypes. We next filtered out the genes spatially enriched for both subtypes, retaining only those uniquely represented in each subtype-specific gene set. This step allowed us to focus our subsequent investigations on pathways more likely unique to either the hypo- or hyperconnectivity subtypes. We next asked whether these gene sets were enriched for autism-relevant transcripts, and if they recapitulated the molecular pathways observed in mouse hypo- and hyperconnectivity subtypes.

Supporting the validity of our cross-species approach, both dysconnectivity subtypes showed significant spatial enrichment for genes differentially expressed in the postmortem cortex of autistic individuals (hypoconnectivity, OR=1.87, p_(FDR)_=10^-3^; hyperconnectivity, OR=1.88, p_(FDR)_=10^-3^, **Figure 5a**, ^13^). Notably, neither subtype showed significant enrichment for genes associated with bipolar disorder, psoriasis, dementia, ADHD, or schizophrenia (**Supplementary Figure 6**).

**Figure 5.**
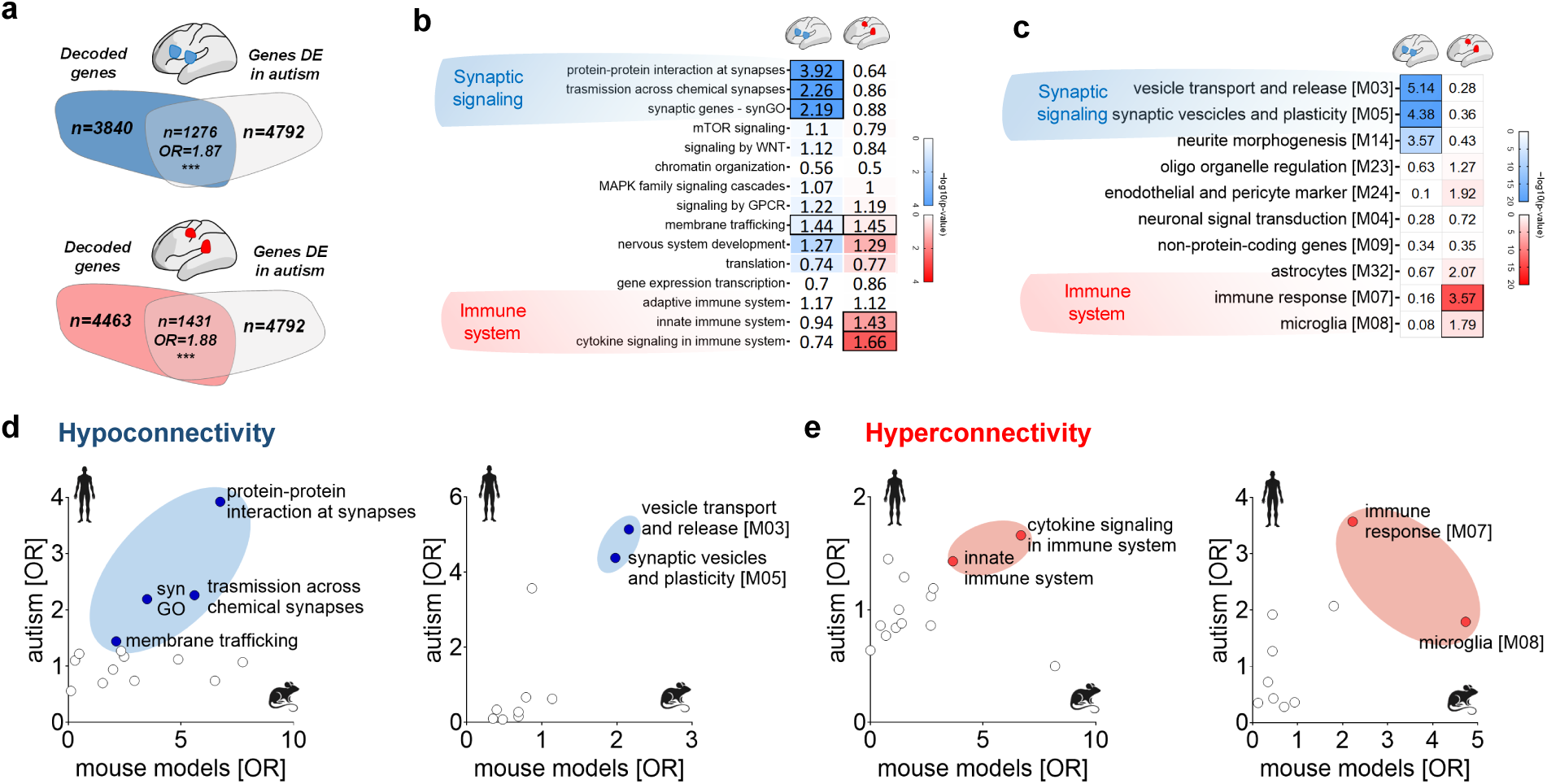
Hypo- and hyperconnectivity subtypes recapitulate synaptic and immune pathways modeled in mice. **a)** Venn diagrams showing enrichment between brain-decoded genes (i.e., genes spatially correlated with dysconnectivity patterns) and differentially expressed genes in autism (Gandal et al., 2022) for both subtypes. Blue/red areas report the number (N) of brain-decoded genes for each subtype. Grey: differentially expressed genes; overlap (i.e. enriched genes) is reported with corresponding odds ratios. ***p(FDR) < 0.001. **b)** Heatmap of enrichments between brain-decoded genes and autism-dysregulated pathways or **c)** modules of genes differentially expressed in autism ^13^. Left columns: hypoconnectivity (blue); right: hyperconnectivity (red). Color intensity indicates enrichment significance (-log p-value). Thick borders mark significant enrichments (q(FDR) < 0.05). **d)** Left: Scatterplot representation of odds ratio (OR) of the molecular ontologies associated with mouse hypoconnectivity (x-axis), versus OR of the same molecular ontologies decoded in autism hypoconnectivity map (y-axis). Right: the same plot is also reported for ORs of genes differentially expressed in autism ^13^. **e)** Left: Scatterplot representation of odds ratio (OR) of the molecular ontologies of mouse hyperconnectivity (x-axis), versus OR of the same molecular ontologies decoded in autism hyperconnectivity map (y-axis). Right: the same plot is also reported for ORs of genes differentially expressed in autism ^13^. Pathways significantly enriched at q(FDR) < 0.05 in both mouse models and autism are highlighted with blue (hypoconnectivity) or red (hyperconnectivity) shading. We report the list of brain decoded genes and the pathways we probed in **Supplementary Table** 3.

Having established that the decoded gene sets were enriched for autism-associated genes, we next investigated their biological functions using pathway-specific gene ontology analysis. This analysis revealed that the two subtypes are associated with distinct biological pathways, mirroring specific molecular dysfunctions observed in the corresponding rodent connectivity subtypes (**Figure 5b-c**). Specifically, brain-decoded genes for the hypoconnectivity subtype showed robust enrichment for multiple synaptic ontologies, such as protein-protein interaction at the synapsis (OR=0.92, p_(FDR)_=10^-6^), transmission across chemical synapsis (OR=2.26, p_(FDR)_=10^-7^), SynGO (OR=2.19, p_(FDR)_=10^-14^), and membrane trafficking (OR=1.44, p_(FDR)_=10^-2^). In contrast, the hyperconnectivity subtype was specifically enriched for immune-related pathways, such as cytokine signaling (OR =1.66, p_(FDR)_=10^-3^) and innate immune system function (OR=1.43, p_(FDR)_=10^-2^, **Figure 5b**), but did not show enrichment for synaptic ontologies (all p_(FDR)_>0.05). Unlike what was observed in our rodent dataset, no enrichment for transcriptional mechanisms was observed in this connectivity subtype. A replication of gene enrichment analysis using gene modules differentially expressed in the autistic brain ^13^ revealed significant synaptic enrichment in the hypoconnectivity subtype (e.g., vesicle transport and release, OR=5.14, p_(FDR)_=10⁻⁵⁶; synaptic vesicles and plasticity, OR=4.38, p_(FDR)_=10⁻²⁶) and immune-related enrichment in the hyperconnectivity subtype (e.g., immune response, OR=3.57, p_(FDR)_=10⁻¹³; reactive microglia, OR=1.79, p_(FDR)_=10⁻², **Figure 5c**). These results suggest cross-species conservation of the pathological pathways associated with the fMRI dysconnectivity subtypes. Similarly, we also found a broad correspondence between the ontologies or gene modules most robustly enriched in rodent dysconnectivity subtypes, and those more prominently enriched in the corresponding human subtypes (**Figure 5d,e**). Collectively, these results corroborate the mechanistic validity of our cross-species decoding and shed light on the molecular dysfunctions underlying two prominent and reproducible brain dysconnectivity subtypes in autism.

## Discussion

Using cross-species functional neuroimaging in large cohorts, we biologically decoded heterogeneous fMRI dysconnectivity patterns related to autism into two reproducible subtypes: one predominantly characterized by whole-brain hypoconnectivity, the other by widespread hyperconnectivity. Notably, gene decoding and enrichment revealed that the identified hypo- and hyperconnectivity subtypes are associated with synaptic dysfunction and immune-related mechanisms, respectively. From a translational standpoint, our results demonstrate the feasibility of using cross-species approaches to empirically decode a widely investigated neuroimaging phenotype in autism into mechanistically dissociable subtypes. Our findings also support the notion that heterogeneous brain dysconnectivity in autism reflects its underlying etiological heterogeneity, thereby providing empirical support for ongoing efforts in neurobiological subtyping of the spectrum ^15,17,19^.

Building upon and expanding our prior rodent fMRI database ^31^, we investigated a broad range of etiologically relevant autism mouse models. Our findings provide compelling evidence that atypical functional connectivity, along with its cross-etiological heterogeneity, is a defining pathophysiological hallmark of autism and is associated with distinct signaling pathways. Leveraging translationally relevant aggregate measures of dysconnectivity, our cross-species analyses uncovered novel biological insights into the molecular underpinnings of autism-relevant fMRI dysconnectivity. The enrichment in multiple synaptic ontologies found in the hypoconnectivity subtype implicates a central role of synaptic dysfunction in altering large scale fMRI connectivity. This phenomenon has been recently investigated at both theoretical and experimental level, revealing a putative direct covarying relationship between excitatory synaptic density, and aggregative fMRI connectivity measures like those used here ^23,25^. The observation of decreased excitatory spine density in multiple mouse lines within the hypoconnectivity subtype, such as *Shank3* ^48^, *Cntnap2* ^49^, *Syn2* ^50^ *16p11.2* ^51^ and *22q11.2* ^25^ broadly supports this hypothesis, and suggests that alterations in synaptic homeostasis (putatively leading to reduced synaptic density) may represent a plausible neurocellular marker for the observed hypoconnectivity.

The robust cross-species association between hyperconnectivity and immune-related signaling is also of great interest, as it suggests that various immune-mediated pathways converge to drive excessive or aberrant functional coupling in the mammalian brain. This may occur through immune-mediated alterations of excitatory or inhibitory function ^52,53^, microstructural white matter abnormalities ^54^, microglial-induced alterations in axonal wiring and synaptic pruning ^55^, as well as immune-related disruption of synaptogenesis ^56^. Importantly, many of these immune-related mechanisms have been shown to directly affect synaptic maturation and homeostasis ^57^. Within this framework, fMRI hyperconnectivity may thus partly reflect an immune-related excess of excitatory synapses. The observation that multiple mouse models within the hyperconnectivity subtype, such as *Cdkl5* ^58^, *Tsc2* ^59^ *Chd8* haploinsufficient mice ^60^, as well as immune-related models such as mice prenatally treated with *Il6* ^61^, and Trem2 deficient mice ^62^, have been reported to exhibit increased synaptic density–at least at early postnatal stages–aligns with this notion and corroborates emerging evidence linking synaptic dysfunction to macroscale fMRI dysconnectivity ^23,25^.

Neuromanipulations studies in rodents ^21^ also enabled us to speculate on the potential neurophysiological determinants of the observed fMRI dysconnectivity. Specifically, using chemogenetics and multielectrode electrophysiological recordings, we recently described an inverse relationship between cortical excitability and fMRI connectivity. Specifically, we found that increased neural firing and cortical excitability may counterintuitively lead to reduced fMRI connectivity ^63,64^, while decreased cortical excitability can lead to increased fMRI connectivity ^65^. According to this model, fMRI hypo- and hyperconnectivity subtypes may thus reflect broadly increased cortical excitability, or excessive inhibition, respectively. The possible coexistence of contrasting excitatory dysfunction within the autistic spectrum would be consistent with the recent identification of electrophysiologically opposed autism subtypes characterized by increased and decreased excitability as measured with EEG ^66^. The hypothesis that the fMRI subtypes we describe here similarly reflect contrasting alterations in excitation-inhibition imbalance warrants empirical testing in both rodents and humans and could have important implications for autism stratification or therapy.

While our cross-species approach primarily focused on two major hypo- and hyperconnectivity subtypes, the relatively coarse partitioning we implemented may have hindered the detection of additional dysconnectivity subtypes, and more nuanced sets of molecular alterations. For example, one set of pathways we were unable to reliably decode in humans is transcriptional dysregulation, which in mice co-clustered with immune dysfunction. It is conceivable that the patterns produced by these two distinct pathways could become dissociable by extending our database to include a larger number of mouse models. Alternatively, transcriptional dysregulation may not be reliably linkable to distinctive pattern of dysconnectivity, owing to the highly stochastic nature of its developmental outcomes ^67^. This hypothesis would be consistent with the phenotypic variability observed upon deletion of the chromatin regulator CHD8 on different genetic backgrounds ^68^.

Notwithstanding the large evolutionary distance between rodents and humans, and our current inability to effectively model common polygenic variants in rodents, our cross-species approach allowed us to successfully decode one fourth of the fMRI scans acquired in individuals with idiopathic and non-syndromic autism. This finding underscores the robust translational potential of our approach and suggests that prevalent autism-relevant pathophysiological alterations can be decoded in human fMRI scans despite key cross-species differences. This result also reinforces previous evidence of substantial overlap between pathways dysregulated in idiopathic autism, and those affected by the rare genetic variants that can be reliably modelled in rodents ^13,27^.

While our current framework does not yet capture the full spectrum of autism-related brain dysconnectivity, our results provide a solid foundation for expanding this approach. The integration of functional neuroimaging in genetically characterized autistic individuals, alongside with future expansions of our rodent database to encompass additional mutations and mechanisms, will be crucial for further refining these insights, with the potential of revealing additional dysconnectivity subtypes beyond the dominant hyper- and hypoconnectivity subtypes we described here. We also note that our findings do not imply that scans lacking subtype classification necessarily exhibit typical or unaltered connectivity, nor do they suggest that putative connectivity changes in this population would be unrelated to underlying etiology. Rather, they suggest that atypical fMRI connectivity in autism exists along a more subtle continuum, often manifesting in ways that are not readily detectable via conventional case-control comparisons. By refining our analytical approaches and expanding our datasets, future studies may capture a broader range of connectivity alterations, further enriching our understanding of the etiological diversity of autism.

The observation of symptom severity differences between the two autism subtypes in the face of their largely opposing functional architecture supports future studies aimed at examining more fine-grained symptom scores. Indeed, deeper and more harmonized phenotyping are critical steps for future translational and clinical subtype validation. Nevertheless, our demonstration that fMRI can encode for complex biological pathways suggests that there is broad scope for fMRI to be used in neurological and psychiatric research, beyond the mere establishment of brain-behavior associations.

In conclusion, our results redefine autism as a neurodevelopmental condition characterized by dissociable neurobiological subtypes; further they provide empirical evidence linking the phenotypic heterogeneity of the autism spectrum to its underlying biological variability. Our work also offers novel insights into the pathobiology of autism, highlighting a role of synaptic dysfunction and immune-related mechanisms in driving autism-related functional dysconnectivity. Finally, from a methodological standpoint our cross-species approach provides an advanced translational framework for a multidimensional, biologically grounded stratification of autism. Our database is openly available to the research community, supporting future investigations into autism-related connectivity alterations.

## Acknowledgments

This work was supported by the Simons Foundation (SFARI 314688, 400101 to A.G., SFARI 982347 to A.G. and M.V.L), the European Research Council (ERC) under the European Union’s Horizon 2020 research and innovation program (#DISCONN; no. 802371 and no. 101125054 #BRAINAMICS to A.G.) and European Union’s Horizon 2020 research and innovation programme under grant agreement No. 845065 (Marie Sklodowska-Curie Global Fellowship - CANSAS to M.P.) The authors also acknowledge support from the NIMHR01MH105506 and R01MH133334 to ADM. We thank Prof. Fabio Benfenati for donating the SynII knockout model. The authors are also grateful to Paige Furano and the CMI staff that collected and curated the data aggregation for the CMI dataset, as well as all the investigators involved with the ABIDE repository creation and all research participants.

## Methods

### Mouse studies

#### Ethical statement

All study procedures were approved by the institutional review board and are in accordance with the ethical standards of the Declaration of Helsinki of 1975, as revised in 2008. All experiments performed at IIT Rovereto were conducted following the Italian Law (DL 27/1992 and DL 26/2014, EU 63/2010, Ministero della Sanità, Roma) and the recommendations in the Guide for the Care and Use of Laboratory Animals of the National Institutes of Health. All experiments performed at ETH Zürich were in accordance with the Swiss federal guidelines for using animals in research and under licensing from the Zürich Cantonal veterinary office. Animal research protocols were also reviewed and approved by the respective animal care committees.

#### Autism mouse models

As detailed in **Table 1**, our collection of fMRI scans in 20 autism-relevant models includes retrospectively aggregated data from published and unpublished experiments across two laboratories. Specifically, twelve mouse models were scanned at IIT Rovereto (Italy) and eight at ETH Zürich (Switzerland). Each study included mice with autism-relevant alterations and wild-type control littermates. Seventeen of these models carried genetic alterations associated with autism; the remaining included a model of environmental autism-risk factor to maternal exposure to interleukin-6 ^61^; a model for TREM2 deficiency ^62^ characterized by microglial defects and autistic-like behavioral phenotype, and inbred BTBR mice ^69^. The BTBR mouse line is characterized by congenital agenesis of the corpus callosum, a neuroanatomical trait associated with high prevalence in autism. A cohort of age-matched C57B6J mice was imaged during the same fMRI session of the BTBR mice and represents the reference control for this line ^70^.

#### Resting-state fMRI

Mice were scanned on a Biospec 70/16 small animal MR system (Bruker BioSpin MRI, Ettlingen, Germany) using controlled sedation ^71,72^. Scans from IIT Rovereto were obtained with a 72 mm birdcage transmit coil and a custom-built saddle-shaped four-element coil for signal reception. Scans from ETH Zürich were obtained with a cryogenic quadrature surface coil (Bruker BioSpin AG, Fällanden, Switzerland). Standard adjustments included calibration of the reference frequency power and the shimming using MapShim (Paravision). Resting state fMRI BOLD time series were acquired using a standard echo planar imaging sequence, as detailed elsewhere ^31^. In all fMRI acquisitions, controlled sedation was obtained using either isoflurane (0.5%) + medetomidine (0.05 mg/kg) for data acquired at ETH Zürich ^73^, or halothane (0.75%) for data acquired at IIT Rovereto ^74–76^, with the exception of OXTR-KO, which were imaged under isoflurane and medetomidine.

### Resting-state fMRI connectivity mapping

Before mapping functional connectivity, we preprocessed the fMRI time series of the autism-related mouse models and wild-type control littermates. Preprocessing encompassed two sequential steps: core preprocessing and denoising. Core preprocessing included the following steps (in brackets the software and function employed). The initial 50 volumes of the time series were removed to allow for T1 and gradient thermal equilibration effects (AFNI 3dTcat). BOLD time series were then despiked (AFNI, 3dDespike), motion-corrected (FSL mcflirt), skull-stripped (FSL bet), and spatially normalized with affine and diffeomorphic registration (ANTS antsRegistration + ANTS antsApplyTransforms) to a skull-stripped reference BOLD template. The denoising pipeline included the following steps: motion traces of six head realignment parameters (three translations + three rotations) and mean ventricular signal (corresponding to the averaged BOLD signal within a reference ventricular mask; FSL fslmeants) were used as nuisance covariates and regressed out from each time course (FSL fsl_regfilt). Then, nuisance regressed time series underwent band-pass filtering to a frequency window of 0.01 - 0.1 Hz (AFNI 3dBandpass) and spatial smoothing with a full width at half maximum of 0.6 mm (AFNI 3dBlurInMask).

fMRI connectivity was quantified on preprocessed timeseries using a voxelwise computation of weighted degree centrality for all mice ^32^. We refer to this parameter here as to “global connectivity” ^25,77^. Global connectivity corresponds to the mean temporal Pearson’s correlation between a given voxel and all the other voxels within the brain ^77^. This approach allowed us to obtain spatially unbiased connectivity mapping without the constraints of pre-imposed anatomical boundaries ^78^. Global connectivity is also amenable to direct cross-species translation ^23,51^. Pearson’s correlation scores were first transformed to z scores using Fisher’s r-to-z transform and then averaged across voxels to yield final global connectivity strength. Finally, differences in global connectivity strength between each autism model and its control littermates were quantified at the voxel level using Cohen’s *d* ^79^. Cohen’s *d* global connectivity maps for each autism-relevant mouse model were then plotted using global histogram analysis as implemented in the R package ggplot. Voxelwise global connectivity mapping was implemented using custom code in Python 3.

### fMRI dysconnectivity subtyping in autism-relevant mouse models

To identify cross-etiological fMRI dysconnectivity patterns we applied cluster analysis to our Cohen’s *d* global connectivity maps. To this end, after vectorizing those statistical maps, we created a connectivity matrix, where rows are autism-relevant mouse models and columns are brain voxels. We then applied agglomerative hierarchical clustering to identify mouse models exhibiting similar Cohen’s *d* global connectivity maps, as implemented in R package heatmap.2. A dendrogram was used to visualize the degree of similarity between autism-relevant mouse models, and to demarcate clusters. Similarity was quantified by using Euclidean distance. To determine the optimal number of clusters, we used NbClust ^80^ and we searched up to ten cluster solutions. The 16 indices calculated by the R function NbClust revealed that k=2 (7/16) was the optimal solution, followed by k=3 (6/16). Considering that a) we searched for dominant brain topographies that could be robustly translated to human brain scans, and b) one of the partitions obtained with k=3 would be composed of only two mouse models, we retained k=2 for subsequent analyses.

To identify brain regions showing prominent cross-etiological fMRI dysconnectivity in each cluster (i.e., subtype), we first thresholded and binarized their global connectivity maps at Cohen’s *d* > 0.8, and then we calculated conjunction maps across autism-relevant mouse models for each subtype. The use of conjunction maps allowed us to identify brain regions that were consistently hypo- or hyper-connected in at least two mouse models belonging to the same subtype. These analyses allowed us to generate two distinct conjunction maps, one showing brain regions predominantly hypoconnected (across mouse lines belonging to the hypoconnectivity subtype), the other one showing brain regions predominantly hyperconnected (across mouse lines belonging to the hyperconnectivity subtype).

### Gene enrichment analysis

To investigate whether different patterns of autism-relevant dysconnectivity reflect dissociable biological pathways, we conducted a gene enrichment analysis. This analysis compares two lists of genes, statistically testing whether a gene set in one list is represented in the other list above chance using a hypergeometric test. Gene enrichment analysis was carried out between a list of autism-relevant genes grouped in the two subtypes of the mouse models (plus their interacting genes), and a list of genes indexing the molecular pathways known to be associated with autism. To this purpose, we created two *in silico* protein-protein mega-interactomes, one for the hypo- and another for the hyper-connected subtype. Each mega-interactome includes all the genes (inferred by the corresponding mouse model) that clustered into either the hypo- or hyperconnectivity subtypes, along with their interacting genes. To search for those interacting genes, we carried out a protein–protein interaction analysis using STRING-DB (Szklarczyk et al., 2019). For monogenic mouse models (e.g. shank3, cntnap2), we used the single mutated or knockout gene as a seed in protein-protein interaction analysis For 16p11.2 ^81^ and 22q11.2 ^82^ we carried out simultaneous seed-based analysis for all the genes included in these two copy number variants (n=27 for 16p11.2 and n=24 for 22q11.2, as modeled in our mice). As an inbred model not relatable to specific genetic alterations, BTBR mice were excluded from this analysis. Overall, these analyses produced a protein-protein interactome of up to 100 genes for each autism etiology. The interactomes of the mouse models within each dysconnectivity subtype were then aggregated to create two mega-interactomes. This analysis produced one mega-interactome of the genes associated with fMRI hypoconnectivity (n=527) and another for those linked to fMRI hyperconnectivity (n=585). We next filtered out genes present in both mega-interactomes (n=70), to retain those uniquely represented in either of the mega-interactomes, resulting in n=457 and n=515 for the hypo- and of the hyperconnectivity subtype, respectively. This step allowed us to focus our subsequent investigations on pathways specifically linked to the two dysconnectivity subtypes.

For each mega-interactome, we next applied a gene enrichment analysis to identify the prevalence of molecular pathways known to be dysregulated in autism as described in ^33^. Molecular pathways included adaptive immune system, chromatin organization, cytokine signaling in immune system, gene expression transcription, innate immune system, MAPK family signaling cascade, membrane trafficking, mTOR signaling, nervous system development, protein-protein interaction at synapses, signaling by GPCR, signaling by WNT, translation, transmission across chemical synapses. The ontologies of the molecular pathways were downloaded from https://reactome.org/. Together with molecular ontologies, we also carried out gene enrichment analysis for a list of manually curated synaptic genes (“SynGo”) ^83^. To probe the robustness of our gene enrichment analysis against interactome size (i.e., the maximum number of interacting genes), we repeated the enrichment using interactomes composed of up to 500 genes. This analysis resulted in n=1793 genes of the interactomes of the hyperconnectivity and n=1526 hyperconnectivity subtypes. After filtering out the n=377 genes shared by the hypo- and hyperconnectivity mega-interactomes, we obtained n=1416 genes uniquely part of the hypoconnectivity subgroup, and n=1149 of the hyperconnectivity subgroup. Enrichment analysis conducted with these larger mega-interactomes resulted in odds ratio highly similar to those obtained with the larger mega-interactomes, thus, ruling out that the size of the interactome could drive results (**Supplementary Figure 1**). Finally, a complementary enrichment analysis was carried out between the meta-interactomes and the genes of the co-expression modules reported to be differentially expressed in postmortem brain tissues of individuals with autism ^13^. Only co-expression modules of known biological function (n=10 ^13^) were considered for this analysis. Enrichments were measured by using hypergeometric statistical testing and quantified with odds ratios (OR) and p-values (FDR-corrected at q<0.05). The gene lists and the interactomes are listed in **Supplementary Table 2.**

### Human studies

#### Sample

To identify autism subtypes informed by the biologically relevant rodent dysconnectivity subtypes, we analyzed resting-state fMRI timeseries from n=940 individuals with idiopathic autism and n=1036 neurotypical controls comprising 38 data collections, 37 selected from the two ABIDE data repositories ^36,37^, and an additional one more recently collected and aggregated at the Child Mind Institute (CMI) ^38,39^. We included data from individuals aged 5 to 30 years old. As illustrated in the data selection flow in **Supplementary Figure 7**, we retained only brain scans of participants with median FD≤0.2 mm ^84^ that successfully underwent co-registration to MNI standard space. Details on each collection, including scan parameters, are at http://fcon_1000.projects.nitrc.org/indi/abide/abide_I.html for ABIDEI, at http://fcon_1000.projects.nitrc.org/indi/abide/abide_II.html for ABIDEII. Details on the CMI data collection are reported in ^38,39^. Demographics and clinical information of the aggregate sample included in our analyses are summarized in **Table 2**.

### Resting state fMRI preprocessing

Before mapping fMRI connectivity, we preprocessed fMRI timeseries with C-PAC v1.6.2^85^. Briefly, we resampled the data to RPI orientation (AFNI 3drefit and AFNI 3dresample) and conducted slice timing correction (AFNI 3dTshift). Next, motion correction (AFNI 3dvolreg) was performed using a two-stage approach in which the timeseries were first co-registered to the temporal mean fMRI image, then a new temporal mean was calculated and used as the target for a second co-registration ^86^. At this second stage, motion parameters based on the Friston 24- parameter model (six motion parameters, their first order derivatives, 12 squared values of these items) were calculated along with framewise displacement. Motion corrected timeseries were then skull-stripped (AFNI 3dAutomask) and mean-based intensity normalized to a factor of 10,000. Then nuisance variable regression was performed with a 24-regressor model of motion and 5 nuisance signals, identified via principal components analysis of signals obtained from white matter, and mean CSF signal (CompCor, ^87^). Brain mask (probability p>0.95) of white matter and CSF were calculated by applying FSL’s fast tool to co-registered structural MRI images ^88^. Functional-to-anatomical co-registration was achieved by boundary-based registration using FSL FLIRT. The residuals of the nuisance variable regression procedure were then processed with bandpass filtering (0.01Hz < f < 0.1Hz) and subsequently smoothed using a 6mm FWHM kernel. Finally, spatial normalization of preprocessed fMRI timeseries to MNI152 space was applied with linear and non-linear registration using ANTs ^89^.

### Resting-state fMRI connectivity mapping

For consistency with mouse data, fMRI connectivity was quantified by applying the same global connectivity mapping described above ^77^. Global connectivity maps were then harmonized across all data collections (n=38) using ComBat ^90^. Before carrying out subtyping, fMRI global connectivity map of autistic participants underwent voxelwise z-scoring normalization against the mean and standard deviation of the fMRI connectivity maps of the neurotypicals to interpret connectivity patterns of the participants with a diagnosis of autism in terms of connectivity increases or decreases relative to neurotypicals.

### Autism subtyping using fMRI connectivity

We used normalized global connectivity maps to discover subtypes of autism based on the cross-species regional approach summarized in **Figure 3**. Specifically, we searched for normalized global connectivity maps of autistic individuals showing patterns of fMRI hypoconnectivity or hyperconnectivity corresponding to those we observed in the mouse subtypes. To establish regional correspondence across species, we selected n=13 cortical and subcortical regions phylogenetically conserved across species. Boundaries of anatomical regions for the mouse brain were defined by using the Allen Mouse Brain Reference Atlas, and those of the human brain were defined by the Harvard Oxford Atlas (**Supplementary Figure 3a**). The more detailed pipeline we used for our cross-species subtyping with fMRI is depicted in **Supplementary Figure 2.** We first used the conjunction maps of each of the two mouse subtypes to generate “dysconnectivity priors” for hypo- and hyperconnectivity. For each conjunction map, we measured the percentage of voxels exhibiting hypo- (or hyperconnectivity) in each of the n=13 regions (Cohen’s *d* > 0.8). By applying k-means clustering to those percentages among the 13 regions, we then separated brain regions with more prominent dysconnectivity from those with less prominent dysconnectivity (hypo or hyper). This strategy led us to the identification of n=5 brain regions with prominent hypoconnectivity, namely anterior and middle cingulate, insula, motor cortex and striatum **(Supplementary Figures 3b**). These regions were aggregated into an aggregated mouse hypoconnectivity mask. For the rodent hyperconnectivity subtype, we identified n=3 brain regions with prominent hyperconnectivity, namely the amygdala, hippocampus, and striatum **(Supplementary Figures 3b**), which we aggregated into a mouse hyperconnectivity mask as above. We next used these two masks to guide the identification of hypo- and hyperconnectivity subtypes in human dataset. Participants on the autism spectrum exhibiting fMRI connectivity lower than one standard deviation in the regions belonging to the hypoconnectivity mask were grouped into the hypoconnectivity subtype (**Supplementary Figure 2**). Similarly, participants with a diagnosis of autism exhibiting fMRI connectivity higher than one standard deviation in the set of regions homologous to the rodent hyperconnectivity mask were grouped into the hyperconnectivity subtype (**Supplementary Figure 2**). To rule out that subtypes could be differentiated by in-scanner head motion, we measured median framewise displacement for each individual across the two subtypes. Group mean comparisons confirmed comparably low levels of head motion across the subtypes (**Supplementary Table 1**).

### Replicability of subtyping

To test the replicability of our fMRI subtypes, we *a priori* split the cohort of autistic individuals into a discovery and a replication dataset and searched for subtypes in both datasets independently using the same method. To create the discovery and replication dataset we used the R function *group_by,* by following the conditions that i) the two datasets must be carefully matched by diagnosis, sex, age and in-scanner head motion, and ii) that one dataset comprises approximately 70% of the sample. The resulting matched datasets comprised 78.5% of the brain scans in the discovery dataset, (N=744 participants with autism, N=807 neurotypicals; N=38 data collections), and 21.5% in the replication dataset (N=196 participants with autism, N=229 neurotypicals, N=38 data collections). Matching between discovery and replication dataset was confirmed by unpaired t-tests for the continuous variables age (t_(1974)_=1.09, p=0.28) and in-scanner head motion (t_(1974)_=0.10, p=0.92), and by chi-square for the categorical variables, such as diagnosis (χ ^2^=0.45, p=0.49) and sex (χ ^2^=1.82, p=0.18). Global connectivity scores of the autistic individuals grouped in the hypo- and hyperconnected subtypes were then z-scored relative to the NT group distribution for the discovery and replication datasets together. Z-scored global connectivity values were then quantified in the 13 evolutionarily conserved regions (**Supplementary Figure 3c,d**).

Replicability was measured with spatial similarity metrics between the fMRI brain maps of the two subtypes identified in the discovery and replication datasets. Specifically, we quantified Dice Coefficients between conjunction maps, obtained after thresholding and binarizing the fMRI connectivity maps of each subtype at t=3.1 (FWER-cluster corrected at p<0.05) in the discovery and replication dataset (hypoconnectivity subtype: **Supplementary Figure 4a**; hyperconnectivity subtype: **Supplementary Figure 4c)**. Replicability was also assessed by computing spatial correlation (Pearson’s *r*) of unthresholded maps (hypoconnectivity subtype: **Supplementary Figure 4b**; hyperconnectivity subtype: **Supplementary Figure 4d)**. Finally, to rule out that the subtyping could be biased by brain scans acquired in a small fraction of laboratories, we repeated the subtyping described above in the aggregated sample after removing the data collections containing the largest number of autistic participants.

### Functional Network Mapping

To investigate the fMRI network atypicalities associated with each subtype, we calculated fMRI connectivity within and between networks by using network-based statistics (NBS) ^45^. To this aim, we first extracted fMRI timeseries from 400 cortical parcels of the Schaffer atlas ^43^ and the 14 subcortical parcels of the Harvard Oxford Atlas ^42^, distributed with FSL. Shaffer’s cortical regions were grouped in the seven canonical resting state functional connectivity networks described in ^44^. We then calculated the ROI-to-ROI functional connectivity matrix for each brain scan by using temporal Pearson’s correlation. We next regressed out the whole-brain mean connectivity signal (i.e., mean correlation of each participant’s 414*414 parcel-to-parcel connectivity matrix) across participants. This step allowed us to assess the network organization of each subtype map beyond the general hyper- and hypoconnectivity characterizing the two autism subtypes. We finally used unpaired t-tests as implemented by the NBS to identify intergroup functional connectivity differences between autistic individuals and neurotypical participants, within each subtype. Statistical results were corrected for multiple comparisons using FWER. We set the univariate threshold to t≥3.1, the network significance at p=0.05 (two-tailed) and we carried out 5000 permutations. This analysis yielded a group-level functional connectivity matrix for each subtype showing the absolute number of ROI-to-ROI fMRI connectivity surviving NBS-statistical thresholding within and between functional networks. To keep into account that functional networks have a different size (and then are formed by a different number of ROIs), we normalized the functional connectivity matrix by the total number of ROI-to-ROI fMRI connectivity, for within and between functional networks. Network-wise fMRI connectivity differences for each subtype were displayed using connectograms as implemented in the R package circlize.

### Behavioral analyses

To investigate whether the autism connectivity subtypes could be differentiated by severity of autistic symptoms, we used the calibrated severity scores (CSS) consistent with the Autism Diagnostic Observation Schedule Second Edition (ADOS-2), a semistructured interactive assessment of social reciprocal communication skills and restricted repetitive behaviors (RRB) ^91,92^ ^93^. CSS scores provide a ten-point severity scale designed to be comparable across different ages and language levels. CSS total scores were available in a subset of the subtyped autism data (n=125 individuals, of which n=38 belonging to the hypoconnectivity subtype, and n=87 to the hyperconnectivity subtype. To similarly assess autism subdomain symptoms, we converted available social affect (SA) and RRB scaled scores into calibrated severity scores based on ^92^ and^94^ **(Supplementary Methods)**. Unpaired t-tests (FDR corrected for multiple comparisons at *q*≤0.05) were used to assess subtype group mean differences in behavioral scores. Secondary analyses explored subtype comparisons in regard to demographics, IQ scores, and rate of psychiatric co-occurrences and psychoactive medication use available **(Supplementary Table 1).**

To examine whether the fMRI maps of the subtypes exhibit spatial topography similar to those of the brain maps associated to cognitive functions, we used the 12 ontology probability maps identified in ^40^. Consistent with ^95^, spatial correspondence was measured as the proportion [%] of voxels by which each neurocognitive map thresholded at *p*<1e-5 overlaps with the mean regressed fMRI maps of each of the subtypes at Cohens’ *d*≥0.2. The resulting spatial correspondences were visualized with radar plots.

### Brain decoding and gene enrichment analyses

To search for an autism-relevant genetic signature in the fMRI connectivity maps of the two autism subtypes, we carried out brain decoding and gene enrichment analysis. The goal of these analyses is to test whether genes exhibiting expression patterns spatially correlated to the fMRI maps of the two subtypes are differentially expressed in autism ^13^. To identify genes that are spatially enriched in the dysconnectivity patterns characteristic of each subtype, we carried out brain decoding with NeuroVault ^96^. The analysis first uses a mixed linear model to compute the similarity between an unthresholded whole-brain fMRI map and spatial patterns of gene expression for each of the six donor’s brains of the Allen Institute Human Brain Gene Expression atlas ^97^. The slopes of these donor-specific linear models encode how similar each gene’s spatial expression pattern is to the fMRI map. Donor-specific slopes were then subjected to a one-sample t-test to identify genes whose spatial expression patterns show consistently high similarity across the donor’s brains to the fMRI map. The resulting list of genes is then thresholded for multiple comparisons, and only the genes with t-statistic values surviving FDR q<0.05 are retained for the following enrichment analysis. To increase specificity of the spatial similarity between gene expression and fMRI brain maps of the two autism subtypes, we retained only genes highly expressed in the brain (n=16796 genes). This analysis resulted in a list of n=3840 genes for the hypoconnectivity subtype, and n=4463 genes for the hyperconnectivity subtype (FDR-corrected, q < 0.05) **(Supplementary Table 3).** We next filtered out genes present in both lists (n=2193), to retain those uniquely represented in either list. This resulted in n=1647 and n=2270 for the hypo- and of the hyperconnectivity subtypes, respectively. This step allowed us to focus our subsequent investigations on pathways uniquely associated with each dysconnectivity subtype. With the lists of fMRI-relevant genes isolated, we then carried out a set of gene enrichment analysis to examine whether our fMRI transcriptomic signatures (i.e., the brain decoded genes) were enriched for specific molecular pathways. The first of these sets comprises genes differentially expressed in autism. This list was obtained by aggregating all the genes belonging to the n=24 coexpression modules reported to be differentially expressed in postmortem brain tissues of autistic individuals ^13^. A second gene set encompassed molecular pathways known to be dysregulated in autism as described in ^33^. This was generated by downloading (from https://reactome.org/) the ontologies of autism-relevant human pathways homologous of those we investigated for the mouse brain. Here we also included a gene enrichment analysis for a list of manually curated synaptic genes (“SynGo”) ^83^. We finally probed enrichment for a third set of genes this time encompassing gene co-expression modules labelled with known biological function (n = 10 ^13^). Enrichment was measured by using hypergeometric statistical testing and quantified with odds ratios (OR) and p-values. Genes surviving correction for multiple comparisons at FDR q < 0.05 were considered statistically significant. A list of brain decoded genes and pathways are reported in **Supplementary Table 3.**

## Code and data availability

The code used for preprocessing mouse fMRI data is available at https://github.com/functional-neuroimaging/fMRI-preprocessing. The code for mapping global connectivity in mice and humans is available at https://github.com/functional-neuroimaging/fMRI-global-local-connectivity. The code for site harmonization is available at https://pypi.org/project/combat/. The code employed for gene enrichment analysis is available at https://github.com/functional-neuroimaging/gene_decoding_and_enrichment. The code of NbClust is available at: https://cran.r-project.org/web/packages/NbClust/NbClust.pdf.

