## Supplementary Figures and Methods for "Biological subtyping of autism via cross-species fMRI"

*[1]Functional Neuroimaging Laboratory, Istituto Italiano di Tecnologia, Center for Neuroscience and Cognitive Systems, CNCS@UNITN, Rovereto, Italy*
*[2]Autism Center, Child Mind Institute, New York, NY, USA*
*[3]IMT School for Advanced Studies, Lucca, Italy*
*[4]Department of Psychiatry, University of Geneva, Switzerland*
*[5]Department of Basic Neurosciences, University of Geneva, Switzerland*
*[6]Center for Mind and Brain Sciences (CIMeC), University of Trento, Rovereto, Italy*
*[7]Brain Research Institute, University of Zurich, Zurich, Switzerland*
*[8]Synaptic Plasticity of Inhibitory Networks, Istituto Italiano di Tecnologia, Genova, Italy*
*[9]Centre for Craniofacial and Regenerative Biology, King's College London, London, UK*
*[10]Department of Clinical and Biomedical Sciences, University of Exeter, Exeter, UK*
*[11]Bloorview Research Institute, Holland Bloorview Kids Rehabilitation Hospital, Toronto, ON, Canada*
*[12]Boston Children's Hospital, Harvard Medical School, Boston, MA, USA*
*[13]Wellcome Centre for Integrative Neuroimaging, FMRIB, Nuffield Department of Clinical Neurosciences, University of Oxford, Oxford, UK*
*[14]Humanitas University, Milan, Italy*
*[15]CNR Institute of Neuroscience c/o Humanitas Clinical and Research Center - IRCCS, Rozzano, Milan, Italy*
*[16]Department of Cellular, Computational and Integrative Biology. University of Trento, Trento, Italy*
*[17]Research Coordination and Support Service, Istituto Superiore di Sanità, Rome, Italy*
*[18]Neural Control of Movement Lab, ETH Zürich, Switzerland*
*[19]Center for Integrative Developing Brain, Child Mind Institute, New York, NY, USA*
*[20]Laboratory for Autism and Neurodevelopmental Disorders, Istituto Italiano di Tecnologia, Center for Neuroscience and Cognitive Systems, Rovereto, Italy*
*[21]Center for the Integrative Developmental Neuroscience, Child Mind Institute, New York, NY, USA*

*+ Current address: IRCCS Ospedale Policlinico San Martino, Genova, Italy*
** These authors contributed equally*

Supplementary Methods

### Measures of autism severity

We assessed autism severity in data with a diagnostic label associated with autism, using ratings consistent with the Autism Diagnostic Observation Schedule-second edition (ADOS-2). We used ADOS as primary measures because of its clinical validity, specificity, and wide use[1,2]. We focused on calibrated severity total scores[3,4]. Further, to assess potential variations in the symptoms contributing to total CSS, we also examined the CSS of the subdomains scales social affect (SA) and restricted repetitive behaviors (RRB)[5]. Calibrated Severity Scores range from 1 to 10, with higher scores indicating more severe autism symptom severity; they allow comparability across ADOS modules which vary by age and language abilities[3-5]. We used these scores to describe the aggregate sample and for group mean comparisons of the identified autism-related brain dysconnectivity subtypes.

For ABIDE data collected and aggregated prior to the publication of the ADOS second edition (ADOS-2)[2], total CSS were computed within each data collection's site based on corresponding algorithms on ADOS-G relevant items per Gotham et al.,[4] (for Modules 2 and 3) and Hus and Lord[3] (for Module 4). As a result, in the present study, total CSS were available for n=549 (58.4%) of the 940 data with an autism diagnostic label included in the aggregated sample. These included all n=63 data from CMI, a subset of the ABIDE I (n=9 data collections: KKI, NYU, UCLA-1, UCLA-2, UM-1, UM-2, USM, Stanford, Yale) and of the ABIDE II (n=11 data collections: GU-1, IP-1, KKI-1, NYU-1, NYU-2, OHSU-1, SDSU-1, SU-2, U-MIA-1, UCD-1, UCLA-1). To examine the role of symptom subdomains, we converted available SA and RRB subscale scaled scores into corresponding CSS based on Hus and Lord [3] guidelines. Social affect (SA) scores were available for n=520 (55.3%) individuals in the autism group. Restricted and repetitive behaviors scores (RRB) were available for n=525 (55.8%) individuals in the autism group.

Supplementary Figures

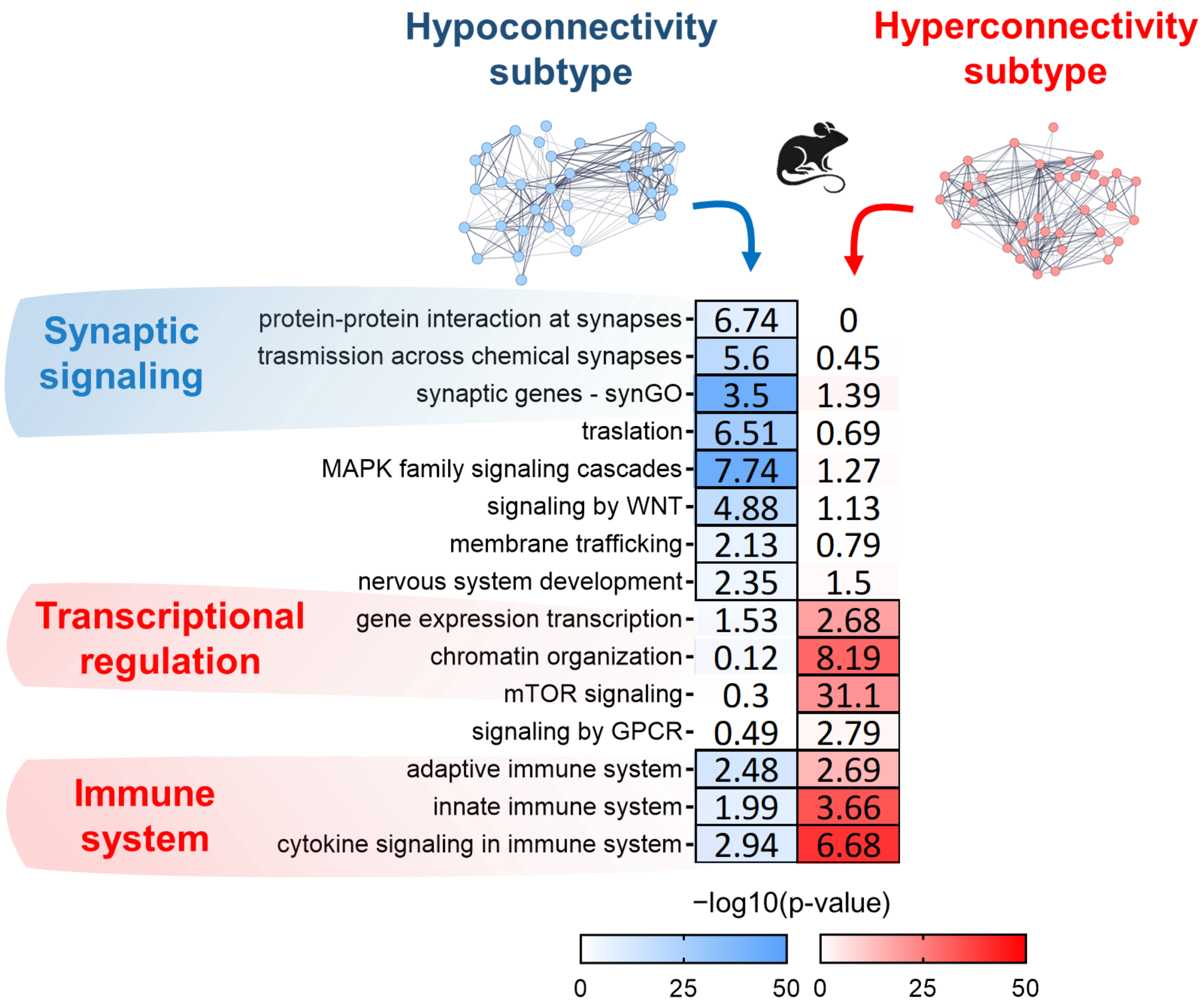


**Supplementary Figure 1. Gene interactome size does not bias mouse subtype pathway enrichment.** Expanded gene interactomes (maximum n=500 interactors per seed gene) recapitulate pathway enrichments obtained using smaller interactomes (n=100, **Figure 2**). Odds ratios for hypoconnectivity subtype are shown in the left column (blue); those for the hyperconnectivity subtype in the right column (red). Cell borders indicate that enrichment is significant at q(FDR) < 0.05.

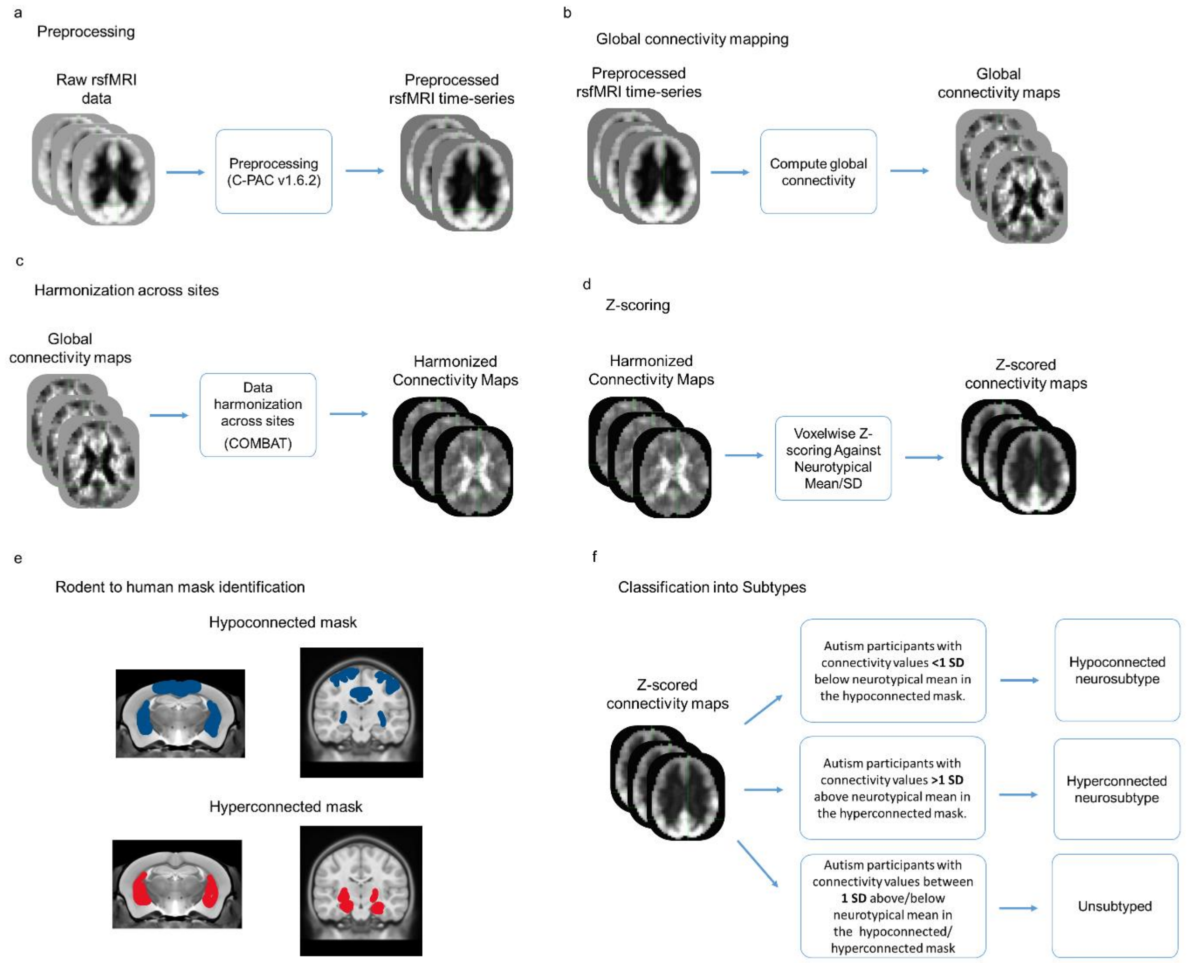


**Supplementary Figure 2. Schematic of fMRI autism subtyping guided by rodent findings. a)** Raw fMRI data from autistic and neurotypical individuals (NTs) were preprocessed using C-PAC v1.6.2. **b)** Preprocessed fMRI timeseries were used to compute global connectivity maps for autism and NTs data. **c)** Global connectivity maps were harmonized across data n=38 data collections with ComBat [6]. **d)** Harmonized global connectivity maps of autism were normalized with Z-scoring against NTs mean and standard deviation. **e)** For each mouse model subtype, we identified a subset of 13 evolutionarily conserved regions (**Supplementary Figure 3**) that best represent predominant dysconnectivity patterns observed across autism-related models. Those regions were combined to generate mouse hypoconnectivity and hyperconnectivity masks. Corresponding hypoconnectivity and hyperconnectivity masks encompassing the same set of brain regions were then created in human data. **f)** Brain scans of autistic individuals were next grouped into subtypes based on the dysconnectivity patterns computed within the masks described above. Autistic individuals with mean global connectivity values in the hypoconnectivity mask more than 1 SD below those of the NTs were assigned to the hypoconnectivity subtype. Those with values more than 1 SD above in the hyperconnectivity mask were assigned to the hyperconnectivity subtype. All the remaining data were considered "unsubtyped".

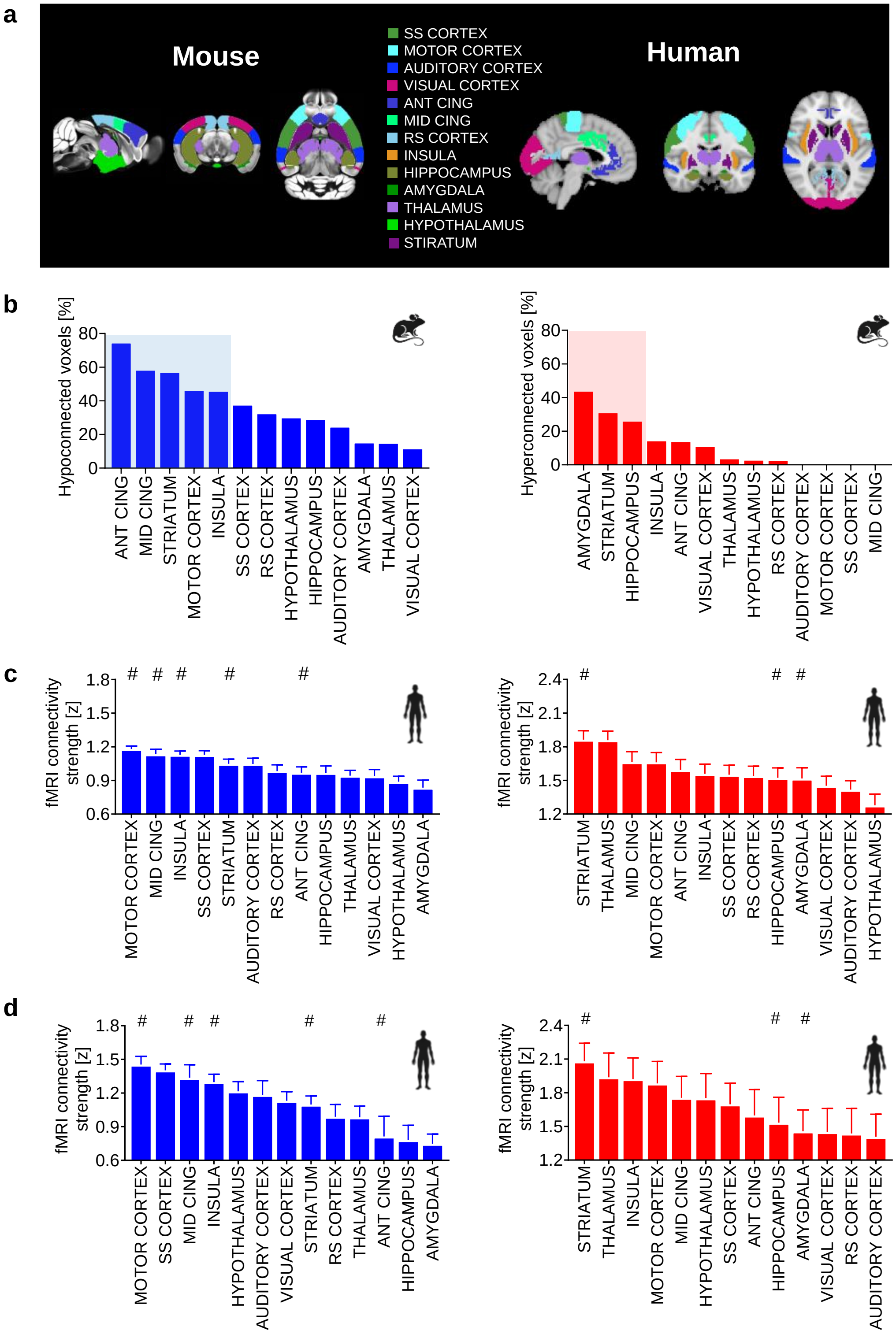


**Supplementary Figure 3. Regional quantification of fMRI connectivity in evolutionarily conserved brain regions in the mouse and human brain. a)** Illustration of the 13 evolutionarily conserved anatomical regions we selected for cross-species extrapolation of fMRI findings. Homologous regions are depicted with the same color. **b)** Regional quantification of the percentage of voxels exhibiting altered connectivity in the mask of the hypo- (left panel) and hyperconnectivity (right panel) rodent subtypes. Bars represent the proportion of voxels exhibiting hypo- (left) or hyperconnectivity (right). Mouse brain regions exhibiting the highest dysconnectivity (shaded in blue or red) were merged together to produce a mask that we later used as a "dysconnectivity prior" for cross-species subtyping. **c)** Regional quantification of atypical connectivity in autism hypo- (left) and hyperconnectivity (right) subtypes in the discovery dataset. **d)** Regional quantification of atypical connectivity of the autism hypo- (left) and hyperconnectivity (right) subtypes in the replication dataset. Error bars indicate mean and SEM. # indicates brain regions homologous to those belonging to the corresponding rodent dysconnectivity prior (i.e. those shaded in b). ANT CINGULATE, anterior cingulate; MID CING, middle cingulate; RS CORTEX, retrosplenial cortex; SS CORTEX, somatosensory cortex.

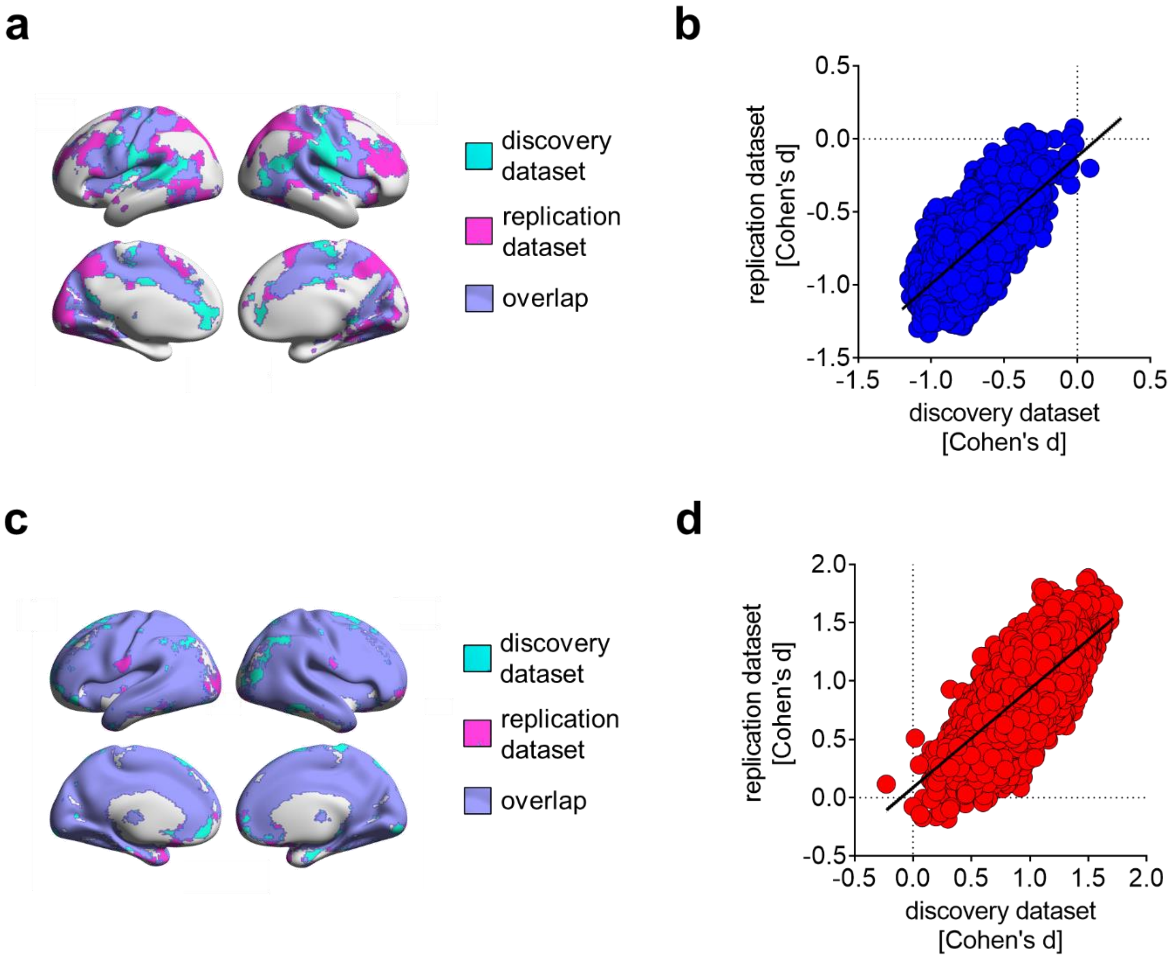


**Supplementary Figure 4. Autism subtypes are replicable. a)** Spatial overlap of hypoconnectivity subtype maps obtained independently in the discovery and replication autism datasets selected *a priori* from the aggregated sample (Dice coefficient=0.74). Light blue indicates regions exhibiting hypoconnectivity uniquely in the discovery dataset. Purple indicates regions exhibiting hypoconnectivity uniquely in the replication dataset. Violet indicates regions exhibiting hypoconnectivity in both datasets (labelled as "overlap"). **b)** Voxelwise spatial correlation between fMRI connectivity maps of discovery and replication datasets in the hypoconnectivity subtype (Pearson's r=0.67). Each circle represents a voxel. **c)** Spatial overlap of hyperconnectivity subtype maps obtained, independently, in the discovery and replication datasets (Dice coefficient=0.96). **d)** Voxelwise spatial correlation between fMRI connectivity maps of discovery and replication datasets in the hyperconnectivity subtype (Pearson's r=0.73). Light blue indicates regions exhibiting hyperconnectivity uniquely in the discovery dataset. Purple indicates regions exhibiting hyperconnectivity uniquely in the replication dataset. Violet indicates regions exhibiting hyperconnectivity in both datasets (labelled as "overlap").

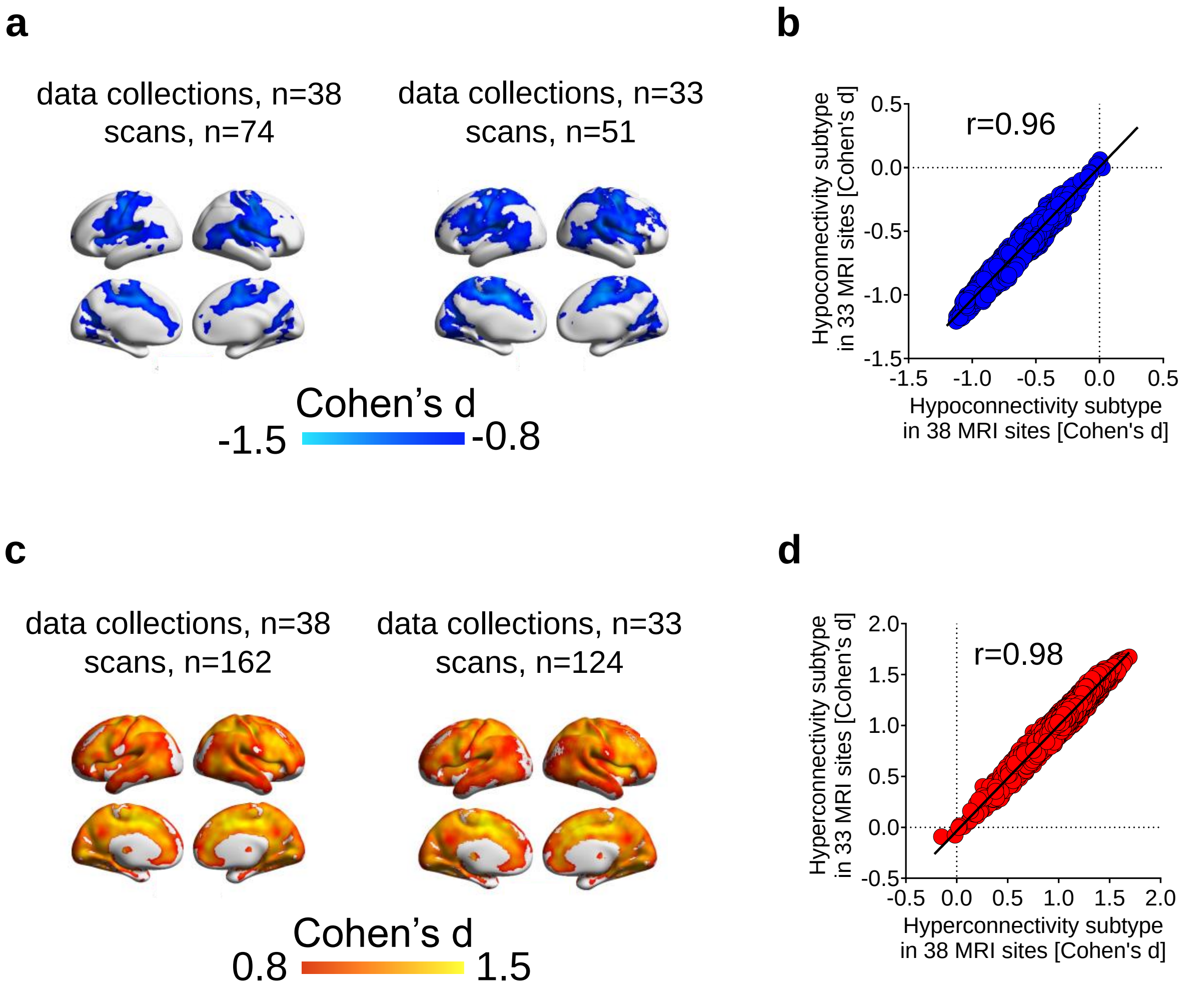


**Supplementary Figure 5. Atypical connectivity in the hypo- and hyperconnectivity subtypes is not driven by the largest autism data collections. a)** Hypoconnectivity subtype maps obtained using all available n=38 data collections (n=74 hypoconnectivity scans, left panel), or upon exclusion of the scans included in the n=5 data collections with the largest autism datasets (n=51 hypoconnectivity scans, right panel). The n=5 data collections are ABIDEI-NYU (n=76 ASD), CMI (n=63 ASD), ABIDEII-NYU-1 (n=47 ASD), ABIDEII-KKI-1 (n=45 ASD) and ABIDEI-USM (n=40 ASD). **b)** Voxelwise spatial correlation between the two hypoconnectivity maps (Pearson's r=0.96). **c)** Hyperconnectivity subtype maps obtained using all available n=38 data collections (n=162 hyperconnectivity scans, left panel), or upon exclusion of the scans included in the n=5 data collections with the largest sample size (n=124 hyperconnectivity scans, right panel). **d)** Voxelwise spatial correlation between the two hyperconnectivity maps (Pearson's r = 0.98).

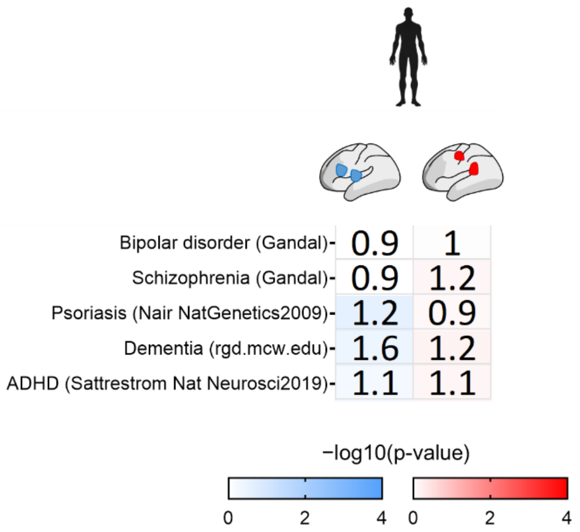


**Supplementary Figure 6. Additional gene enrichment analyses.** The two subtypes did not show significant enrichment for genes associated with bipolar disorders (hypoconnectivity, OR=0.9, $p_{(FDR)}$ = 0.96; hyperconnectivity, OR=1, $p_{(FDR)}$ = 0.93), schizophrenia (hypoconnectivity, OR=0.9, $p_{(FDR)}$ = 0.99; hyperconnectivity, OR=1.2, $p_{(FDR)}$ = 0.67), psoriasis (hypoconnectivity, OR=1.2, $p_{(FDR)}$ = 0.25; hyperconnectivity, OR=0.9, $p_{(FDR)}$ = 0.82), dementia (hypoconnectivity, OR=1.6, $p_{(FDR)}$ = 0.36; hyperconnectivity, OR=1.2, $p_{(FDR)}$ = 0.58), or ADHD (hypoconnectivity, OR=1.1, $p_{(FDR)}$ = 0.51; hyperconnectivity, OR=1.1, $p_{(FDR)}$ = 0.66). ADHD: attention deficit and hyperactivity disorder. These gene lists are reported in **Supplementary Table 4.**

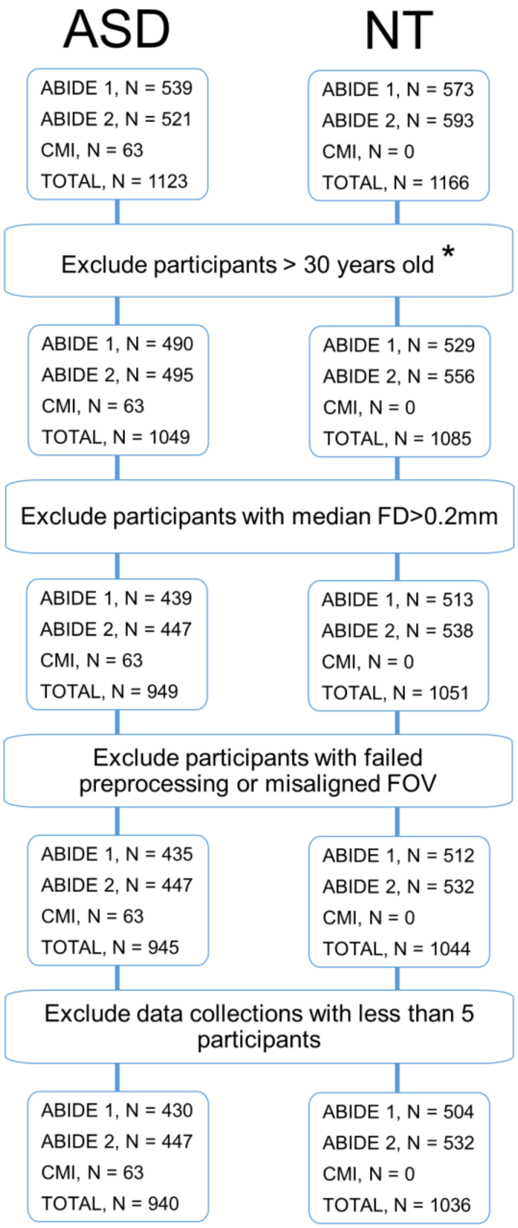


**Supplementary Figure 7. Selection flowchart for the human sample.** The flowchart illustrates the selection process resulting in the cohort of individuals with a diagnostic label of autism spectrum disorder (N=940 ASD) and neurotypical controls (N=1036 NTs) included in this study. At each flowchart step, we report the number of individual data retained from ABIDEI, ABIDEII repositories and within the collections from CMI as total and by diagnostic group. As a result of this selection process, we examined N=1976 brain scans across 23 data collection MRI sites and 38 data collections, including 18 collections from ABIDEI, 19 collections from ABIDEII and one from CMI. Specifically, ABIDEI included: Caltech (ASD n=14; NT n=13), KKI (ASD n=20; NT n=33), Leuven-1 (ASD n=13; NT n=15), Leuven-2 (ASD n=15, NT n=20), MaxMun (ASD n=11; NT n=20), NYU (ASD n=76; NT n=101), OHSU (ASD n=13, NT n=15), Olin (ASD n=18; NT n=15), Pitt (ASD n=22, NT n=23), SDSU (ASD n=13, NT n=22), Stanford (ASD n=18; NT n=20), Trinity (ASD n=23, NT n=24), UCLA-1 (ASD n=39, NT n=31), UCLA-2 (ASD n=9, NT n=13), UM-1 (ASD n=42, NT n=54), UM-2 (ASD n=13, NT n=21), USM (ASD n=43, NT n=36), Yale (ASD n=28, NT n=28). ABIDEII included: BNI-1 (ASD n=12, NT n=10), EMC-1 (ASD n=22, NT n=24), ETH-1 (ASD n=8, NT n=22), GU-1 (ASD n=43, NT n=53), IP-1 (ASD n=19, NT n=23), IU-1 (ASD n=16, NT n=17), KKI-1 (ASD n=45, NT n=148), KUL-3 (ASD n=23, NT n=0), NYU-1 (ASD n=47, NT n=30), NYU-2 (ASD n=27, NT n=0), OHSU-1 (ASD n=36, NT n=55), ONRC-2 (ASD n=22, NT n=30), SDSU-1 (ASD n=33, NT n=24), SU-2 (ASD n=20, NT n=20), TCD-1 (ASD n=16, NT n=19), UCD-1 (ASD n=17, NT n=14), UCLA-1 (ASD n=14, NT n=16), U-Mia-1 (ASD n=13, NT n=15), USM-1 (ASD n=14, NT n=12). CMI included ASD n=63 and NT n=0. *Participants >30 years of age were excluded from the analysis as they correspond to less than 5% of the ABIDE repository. FOV, field of view
