## Supplementary Table 1 for "Biological subtyping of autism via cross-species fMRI"

|  | **Hypoconnectivity autism subtype** | **Hyperconnectivity autism subtype** | **Group comparisons, statistics and p-values** |
| --- | --- | --- | --- |
| **Sample size**  # | 74 | 162 | **-** |
| **Data collections***  # | 28 | 38 | - |
| **Sex**  # M, F | 59,15 | 139, 23 | χ _(1)_^2^ =1.38,   p=0.24,  p_(FDR)_=0.61 |
| **Age**  years | 14.5 (5.4)  [7.2-28.0] | 13.7 (5.2)  [5.6-28.1] | t_(234)_=1.05,  p=0.29,  p_(FDR)_=0.61 |
| **Full Scale IQ**** standard scores | 106.9 (18.9)  [72-148] | 104.5 (16.3)  [61-142] | t_(199)_=0.94,  p=0.35,  p_(FDR)_=0.61 |
| **Verbal IQ*****  standard scores | 107.7 (18.5)  [69-145] | 105.3 (16.7)  [66-149] | t_(175)_ = 0.86,  p = 0.38,  p_(FDR)_=0.61 |
| **Non-Verbal IQ******  standard scores | 104.8 (19.7)  [71-157] | 103.4 (16.5)  [59-146] | t_(182)_ = 0.48,  p = 0.63,  p_(FDR)_=0.63 |
| **median FD**  mm | 0.061 (0.04)  [0.02-0.18] | 0.064 (0.04)  [0.01-0.18] | t_(234)_ = 0.62,  p = 0.53,  p_(FDR)_=0.63 |
| **Psychiatric comorbidity rate^**  n(%) | 12 (48) | 34 (50) | χ _(1)_^2^=0.024,  p=0.62,  p_(FDR)_=0.63 |
| **Psychoactive medication use ^^**  n(%) | 47(34) | 22(39) | χ _(1)_^2^ = 4.04,  p = 0.04,  p_(FDR)_=0.32 |

**Supplementary Table 1. Demographics and characteristics of the fMRI hypo- and hyperconnectivity autism subtypes.** For continuous variables, group mean, and standard deviations are reported in parentheses and minima and maxima are reported in brackets. * The data aggregate of the hypoconnectivity subtype included participants with ASD from all data collections but ABIDEII-ETH-1, ABIDEII-IP-1, ABIDEII-NYU-1, ABIDEII-NYU-2, ABIDEII-UCD-1, ABIDEII-UCLA-1, ABIDEI-MaxMun, ABIDEI-OHSU, ABIDEI-SDSU, ABIDEI-UCLA-2. The data aggregate of the hyperconnectivity subtype included participants with ASD from all n=38 data collections. ** Full Scale IQ was available for n=143 individuals included in the hypoconnectivity subtype and n=58 individuals in the hyperconnectivity subtype. *** Verbal IQ was available for n=124 individuals in the hypoconnectivity subtype and n=53 individuals in the hyperconnectivity subtype. **** Non-Verbal IQ was available for n=128 individuals in the hypoconnectivity subtype and n=56 individuals in the hyperconnectivity subtype. ^ Number and percentage of individuals with one or more psychiatric diagnosis cooccurring with autism. Comorbidity was assessed in a subset of data; specifically, it was assessed in n=27 individuals in the hypoconnectivity subtype and n=68 individuals in the hyperconnectivity subtype. ^^ Number and percentage of individuals using psychoactive medications among the subset of data with information on their use (i.e., n=56 individuals in the hypoconnectivity subtype and n=137 individuals in the hyperconnectivity subtype).  M, males; F, females; FD, framewise displacement; χ2, chi-square statistics; t, unpaired t-test statistics.
